# Eosinophil deficiency promotes aberrant repair and adverse remodelling following acute myocardial infarction

**DOI:** 10.1101/750133

**Authors:** Iqbal S Toor, Dominik Rückerl, Iris Mair, Rob Ainsworth, Marco Meloni, Ana-Mishel Spiroski, Cecile Benezech, Jennifer M Felton, Adrian Thomson, Andrea Caporali, Thomas Keeble, Kare H Tang, Adriano G Rossi, David E Newby, Judith E Allen, Gillian A Gray

## Abstract

**Background:** Eosinophil count predicts outcome following myocardial infarction (MI) and eosinophils regulate tissue repair and regeneration in extra-cardiac settings.

**Objectives:** To investigate the role of eosinophils in regulating inflammation, repair and remodelling following MI.

**Methods:** Blood eosinophil count was assessed in 732 patients undergoing primary percutaneous coronary intervention for ST-segment elevation MI (STEMI). Experimental MI was induced in wild-type (WT) and eosinophil-depleted mice (ΔdblGATA or anti-Siglec F antibody treated). Cardiac function was characterised by high-resolution ultrasound and immune cell infiltration by flow cytometry of infarct digests.

**Results:** Blood eosinophil count declined in the hours following STEMI in patients and following MI induction in mice. Eosinophils were subsequently identified in the myocardium of patients and mice. Genetic eosinophil depletion in mice increased LV dilatation (end-systolic area: 29.0±2.2cm^2^ v 21.6±1.6cm^2^; p=0.02) and reduced ejection fraction (22.0±3.6% v 34.3±4.0% in p=0.04,) in ΔdblGATA v WT following MI (n=8-9/group), an outcome reproduced by pharmacological depletion. ΔdblGATA mice had increased scar size with disrupted collagen deposition and altered expression of the collagen cross-linking genes *plod2* and *lox*. CD206^+^pro-repair macrophages were less prevalent in the infarct zone of ΔdblGATA mice (27.0±2.3% v WT: 39.2.4±3.5%; p=0.01, n=9-11/group) but were restored by replenishment with bone marrow-derived eosinophils. Anti-inflammatory cytokine concentrations were reduced in ΔdblGATA mice and IL-4 complex administration 24h after MI rescued adverse remodelling.

**Conclusions:** Eosinophils are recruited to the heart following MI and are required for effective repair and to prevent adverse remodelling. IL-4 therapy has potential to limit detrimental outcomes when eosinophil availability is low.

**HIGHLIGHTS:** A drop in eosinophil blood count is associated with recruitment of eosinophils to the heart during repair following clinical and experimental myocardial infarction.

Genetic & pharmacological eosinophil depletion leads to increased adverse remodelling in experimental MI.

Eosinophils are required for aquisition of an anti-inflammatory macrophage phenotype and a shift to resolution of inflammation during infarct repair.

Interleukin-4 therapy is able to rescue the adverse remodelling phenotype in conditions of eosinophil deficiency.

**Condensed abstract:** In ST-segment elevation myocardial infarction (STEMI) of both patients and mice, there was a decline in blood eosinophil count, with activated eosinophils recruited to the infarct zone. Eosinophil deficiency resulted in attenuated anti-inflammatory pro-repair macrophage polarization, enhanced myocardial inflammation, increased scar size and deterioration of myocardial structure and function. Adverse cardiac remodelling in the setting of eosinophil deficiency was prevented by IL-4 therapy.

## INTRODUCTION

Myocardial infarction (MI) occurs most commonly following acute thrombotic occlusion of a coronary artery, and triggers an acute inflammatory response. Within hours, neutrophils are recruited to the infarcted myocardium followed by infiltration of pro-inflammatory monocytes.^1^ Acquisition of a pro-resolution proliferative macrophage phenotype is critical to successful infarct repair.^2^ Failure to expand the number of CD206+ pro-resolution macrophages following MI is associated with increased cardiac rupture and adverse cardiac remodelling due to disrupted collagen fibril formation during infarct healing.^2^ Interventions that polarize macrophages towards this phenotype, including interleukin-4 (IL-4),^2,3^ improve infarct healing, but the endogenous mechanisms that regulate repair are not well understood.

Macrophage phenotype can be determined by environmental factors and importantly by cells of the innate and adaptive immune system, including eosinophils. Tissue eosinophilia is commonly associated with helminth infection, allergy and gastrointestinal disorders. In these settings, eosinophils are effector cells with pro-inflammatory and destructive capabilities through the secretion of cytotoxic granules, cationic proteins and proteolytic enzymes.^4^ However, eosinophils also express a number of immuno-modulatory cytokines and lipid mediators implicated in the resolution of inflammation.^4^ Previous studies have linked peripheral blood eosinophil count post-MI to short-term risk of mortality in low-intermediate risk patients^5^ and high risk patients^6,7^ following myocardial infarction. However, whether eosinophils have any role in repair of the adult mammalian heart is so far unknown.

The primary goal of the present study was to address the role of eosinophils in repair of the heart following MI in the setting of patients with acute ST-segment elevation MI (STEMI) and in an experimental model of MI in mice with genetic (ΔdblGATA^8^) or pharmacologic depletion of eosinophils.

## METHODS

### Patient selection with ST-segment elevation MI

All patients included in the STEMI registry were ≥18 years old, had chest pain of <12 hours duration, and persistent ST-segment elevation in at least two contiguous leads of the electrocardiogram.^9^ Patients with a history of stable angina undergoing elective percutaneous coronary intervention were used as the control group. This study was reviewed and approved by the Essex Cardiothoracic Centre Clinical Governance body.

### Animal experiments

12 to 14-week-old WT male BALB/c mice and C57Bl/6 mice were purchased from Harlan Laboratories (UK). ΔdblGATA mice on a BALB/c background^8,10^ were bred and maintained at the University of Edinburgh. All animal experiments were approved by the University of Edinburgh Animal Welfare and Ethical Review Board & the UK Home Office.

### Infarct model

MI was induced by permanent ligation of the left coronary artery in isofluorane anaesthetized mice as previously described.^11^ 24h after surgery, a tail blood sample was collected for assay of troponin I (Life Diagnostics High Sensitivity Mouse Cardiac Troponin-I ELISA kit) to assess the extent of myocardial injury.

### Anti-Siglec-F antiserum depletion of eosinophils

In the eosinophil depletion study, C57BL/6 mice received 100 μL of either sheep anti-Siglec-F polyclonal antiserum or sheep pre-immune serum (both gifted by Professor Paul Crocker, University of Dundee) via intra-peritoneal injection at 1 day before and 3 days after MI.

### Rescue experiments in ΔdblGATA mice

Bone marrow-derived eosinophils were adoptively transferred by intra-peritoneal injection into ΔdblGATA mice at 1 day before and 3 days after MI. Purity of greater than 90% for eosinophils was confirmed prior to intra-peritoneal injection by flow cytometric analysis.

For IL-4 replenishment, 5 μg of recombinant murine IL-4 (Peprotech) complexed to 25 μg anti-IL-4 antibody (clone 11B11; BioXcell) in 100 μL sterile Phosphate buffered saline (PBS) or 100 μL sterile PBS was administered to ΔdblGATA mice via intra-peritoneal injection at Day 1 post-MI.

### Cardiac imaging

Left ventricular structure and function was assessed using high-resolution ultrasound (VisualSonics Vevo 770, Toronto, Canada), under light isoflurane anaesthesia, ensuring that the heart rate was maintained >500 beats/min.

### Histopathology

Eosinophils were detected in samples of paraffin embedded infarcted human myocardium, obtained from the Edinburgh Brain & Tissue Bank (Online Table 2), using eosinophil peroxidase antibody (supplied by Elizabeth Jacobsen, Mayo Clinic, Scottsdale, Arizona).

**Table 1:** Baseline characteristics of stable angina and STEMI patients.

|  | Stable Angina patients<br>N=307 | STEMI patients<br>N=732 | p-value |
| --- | --- | --- | --- |
| Age (yrs) | 65.8 ± 0.5 | 64.6 ± 0.5 | 0.37 |
| Male | 237 (77%) | 547 (75%) | 0.43 |
| Diabetes Mellitus | 79 (26%) | 87 (12%) | <0.001 |
| Hypertension | 189 (62%) | 282 (39%) | <0.001 |
| History of smoking | 153 (50%) | 374 (51%) | 0.63 |
| Hypercholesterolaemia | 250 (81%) | 182 (25%) | <0.001 |
| CVA | 10 (3%) | 36 (5%) | 0.32 |
| Previous MI | 96 (31%) | 92 (13%) | <0.001 |
| Previous CABG | 26 (67%) | 13 (33%) | <0.001 |
| Previous PCI | 113 (65%) | 61 (35%) | <0.001 |
| PVD | 8 (3%) | 14 (2%) | 0.48 |
| Admission Creatinine (mmol/L) | 90 ± 1 | 87 ± 1 | 0.25 |

CVA = cerebrovascular accident, CABG = coronary artery bypass surgery, MI = myocardial infarction, PCI = percutaneous coronary intervention, PVD = peripheral vascular disease, STEMI = ST-segment Elevation Myocardial Infarction.

**Table 2:**
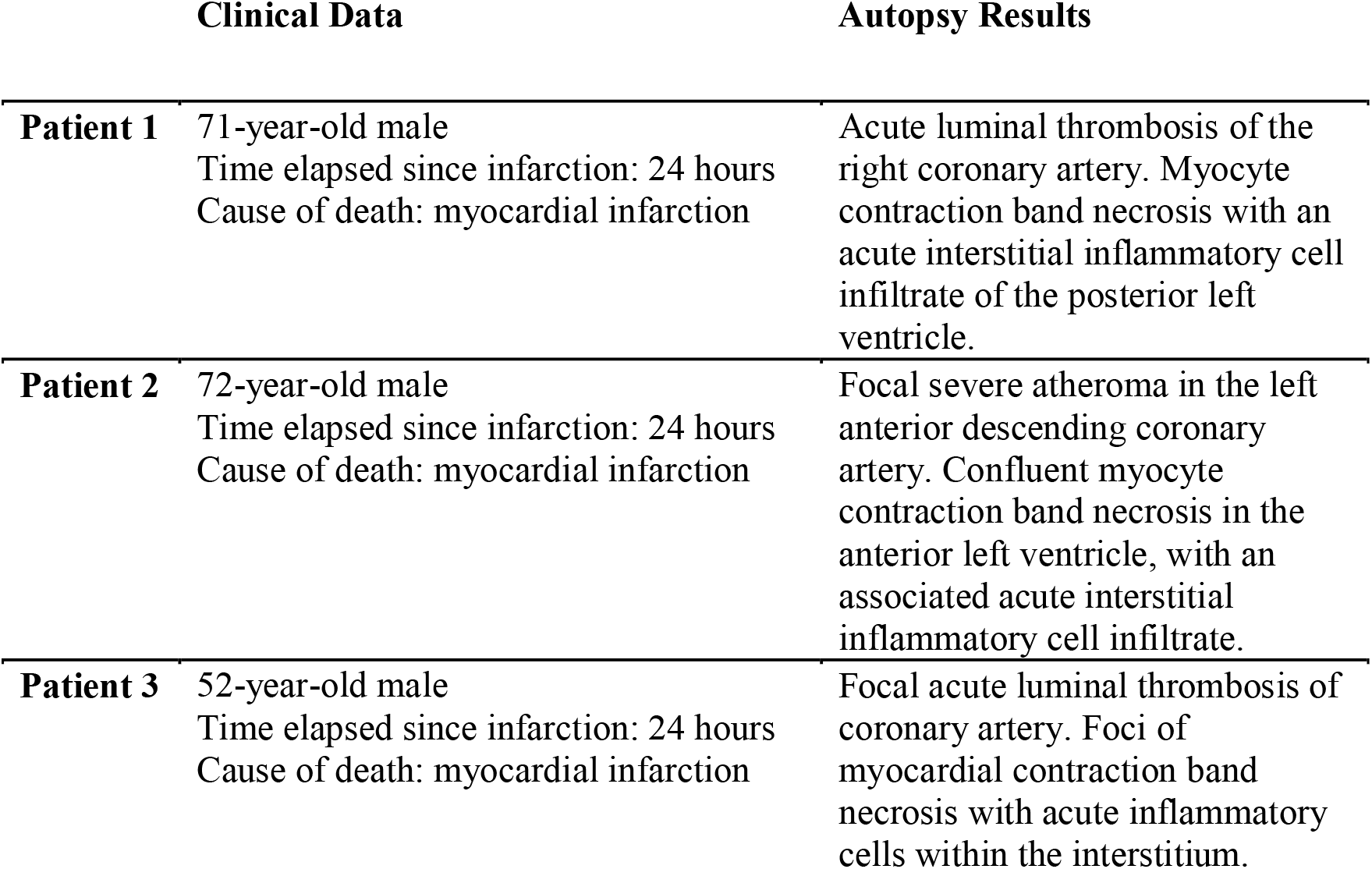
Clinical data and autopsy results for donors of post-MI myocardial tissue used in immunohistochemistry.

Paraffin embedded mouse heart sections were stained with Masson’s trichrome (infarct size) and picrosirius red (polarized light to collagen fibre status using polarizesd light). Angiogenesis was determined by detection of CD31 immunopositive vessels (rabbit antimouse CD31, 1:50; Abcam).

### Transmission Electron Microscopy

Mouse hearts were fixed in 1% Osmium Tetroxide in 0.1M Sodium Cacodylate X3) and embedded in TAAB 812 resin. Ultrathin sections, 60nm thick were stained in Uranyl Acetate and Lead Citrate then viewed in a JEOL JEM-1400 Plus TEM. Eosinophils were identified by their typical morphology with crystalloid granules.

### Flow cytometry

Immunofluorescence staining was performed on tail blood samples and single cell suspensions from heart digests, spleens and peritoneal cells. Flow cytometric analysis was performed on an LSR II instrument (BD Biosciences) and analyzed using FlowJo software (Tree Star). Infarct zone CD45^+^CD11b^+^Ly6G^-^F4/80^+^ macrophages were sorted by FACS using a FACS ARIA II flow cytometer (BD Biosciences). Results for the heart digests are expressed as cell number per infarct zone or remote zone, and total counts were calculated for spleens and the peripheral blood.

### RNA extraction and real-time quantitative PCR

RNA was extracted from infarct zone tissue (RNeasy^®^ Mini Kit, QIAGEN) and reverse transcribed to cDNA with the QuantiTect Reverse Transcription Kit. Real-time quantitative PCR was performed using TAQman^®^ gene expression assays (Online Table 8). mRNA expression levels were normalized for Rpl32 (house keeping gene) expression and are presented as fold changes. Genes were selected based on prior microarray data from the lab (unpublished) that identified them as being significantly modified during the early stages of scar formation, between 3 and 7 days following MI.

**Table 3:**
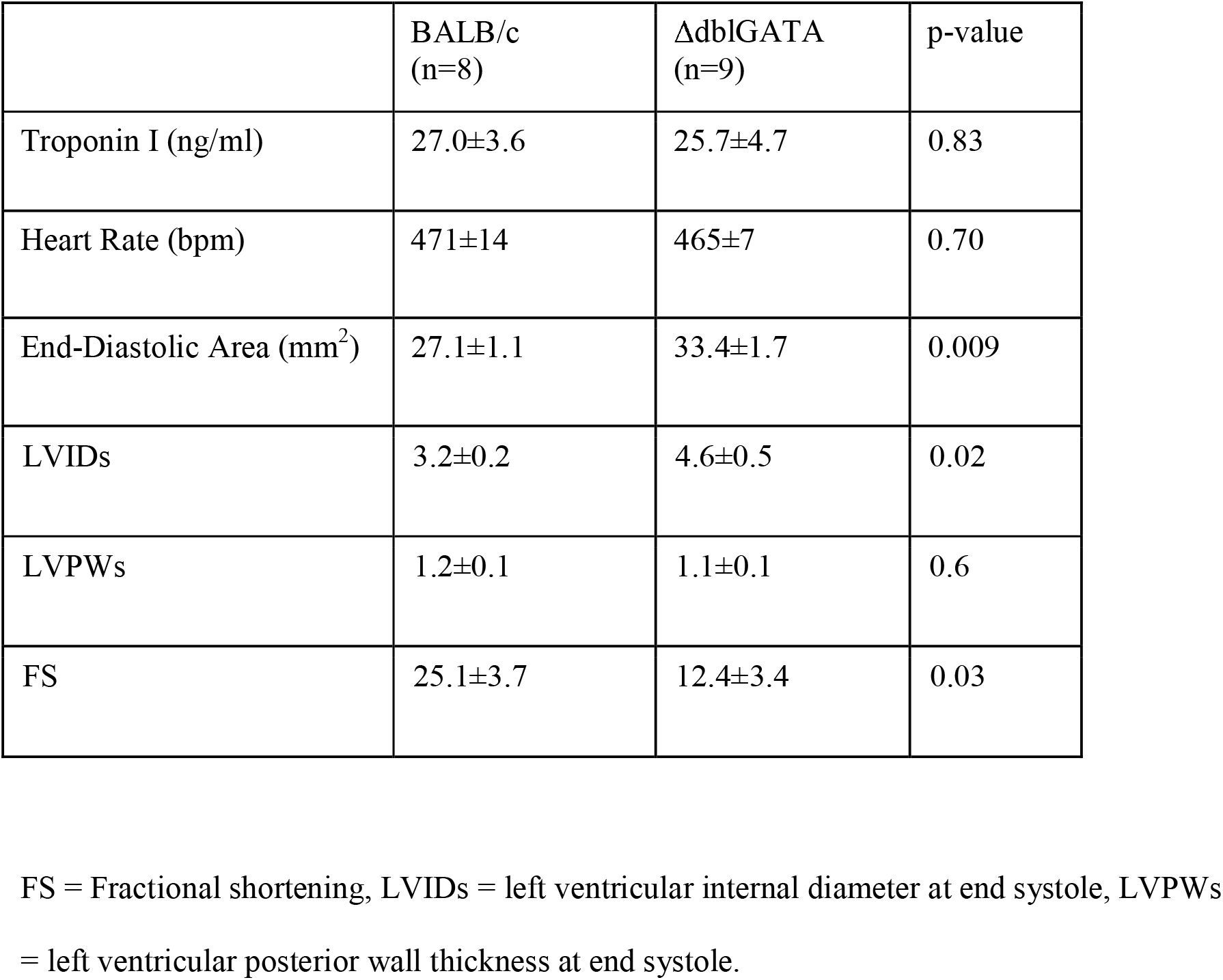
Cardiac function at baseline in WT BALB/c mice and ΔdblGATA mice.

| | BALB/c<br>(n=8) | ΔdblGATA<br>(n=9) | p-value |
| --- | --- | --- | --- |
| Troponin I (ng/ml) | 27.0±3.6 | 25.7±4.7 | 0.83 |
| Heart Rate (bpm) | 471±14 | 465±7 | 0.70 |
| End-Diastolic Area (mm<sup>2</sup>) | 27.1±1.1 | 33.4±1.7 | 0.009 |
| LVIDs | 3.2±0.2 | 4.6±0.5 | 0.02 |
| LVPWs | 1.2±0.1 | 1.1±0.1 | 0.6 |
| FS | 25.1±3.7 | 12.4±3.4 | 0.03 |

FS = Fractional shortening, LVIDs = left ventricular internal diameter at end systole, LVPWs = left ventricular posterior wall thickness at end systole.

**Table 4:** Cardiac injury, structure & heart rate at Day 7 following MI in BALB/c and ΔdblGATA mice.

|  | **BALB/c**<br>**(n=8)** | **ΔdblGATA**<br>**(n=9)** | **p-value** |
| --- | --- | --- | --- |
| **Troponin I (ng/ml)** | **27.0±3.6** | **25.7±4.7** | **0.83** |
| **Heart Rate (bpm)** | **471±14** | **465±7** | **0.70** |
| **End-Diastolic Area**<br>**(mm<sup>2</sup>)** | **27.1±1.1** | **33.4±1.7** | **0.009** |
| **LVIDs** | **3.2±0.2** | **4.6±0.5** | **0.02** |
| **LVPWs** | **1.2±0.1** | **1.1±0.1** | **0.6** |
| **FS** | **25.1±3.7** | **12.4±3.4** | **0.03** |

FS = Fractional shortening, LVIDs = left ventricular internal diameter at end systole, LVPWs = left ventricular posterior wall thickness at end systole, MI = myocardial infarction.

**Table 5:** Cardiac function at Day 7 following MI in C57BL/6 mice injected i.p. with preimmune serum or anti-Siglec-F anti-serum on 1 day before and Day 3 post-MI.

|  | Pre-immune serum<br>(n=6) | Anti-Siglec-F anti-serum<br>(n=6) | p-value |
| --- | --- | --- | --- |
| Troponin I (ng/ml) | 21.9±4.2 | 20.9±3.2 | 0.49 |
| Heart Rate (bpm) | 552±13 | 569±5 | 0.24 |
| End-Diastolic Area (mm<sup>2</sup>) | 26.9±1.4 | 31.9±1.0 | 0.009 |
| LVIDs | 3.0±0.2 | 3.9±0.2 | 0.02 |
| LVPWs | 1.1±0.1 | 1.1±0.1 | 0.6 |
| FS | 28.4±2.0 | 18.9±2.5 | 0.009 |

FS = Fractional shortening, i.p. = intra-peritoneal, LVIDs = left ventricular internal diameter at end systole, LVPWs = left ventricular posterior wall thickness at end systole, MI = myocardial infarction.

**Table 6:**
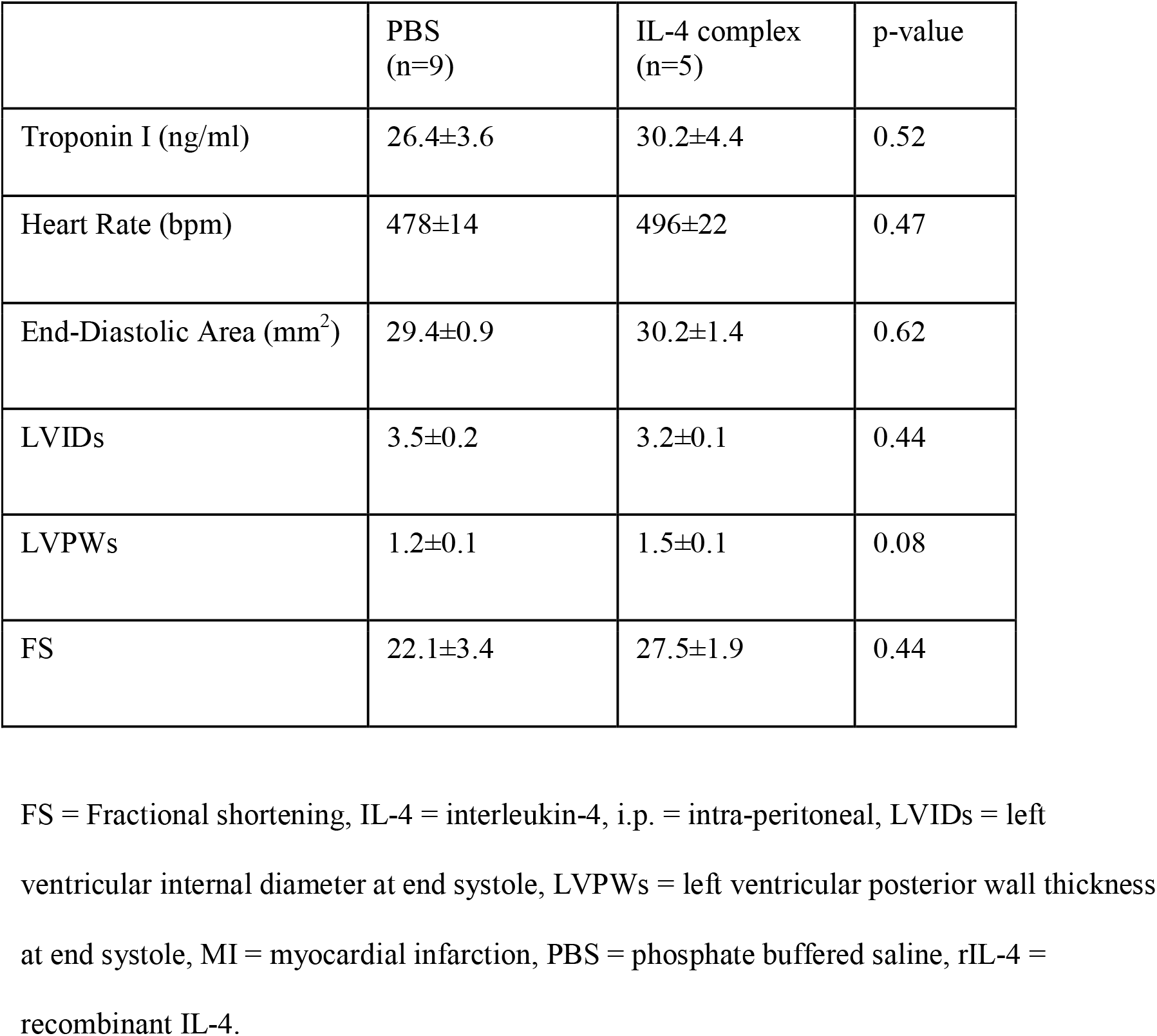
Cardiac function at Day 7 following MI in BALB/c mice injected i.p. with PBS or IL-4 complex containing 5μg rIL-4 on Day 1 post-MI.

|  | PBS<br>(n=9) | IL-4 complex<br>(n=5) | p-value |
| --- | --- | --- | --- |
| Troponin I (ng/ml) | 26.4±3.6 | 30.2±4.4 | 0.52 |
| Heart Rate (bpm) | 478±14 | 496±22 | 0.47 |
| End-Diastolic Area (mm<sup>2</sup>) | 29.4±0.9 | 30.2±1.4 | 0.62 |
| LVIDs | 3.5±0.2 | 3.2±0.1 | 0.44 |
| LVPWs | 1.2±0.1 | 1.5±0.1 | 0.08 |
| FS | 22.1±3.4 | 27.5±1.9 | 0.44 |

FS = Fractional shortening, IL-4 = interleukin-4, i.p. = intra-peritoneal, LVIDs = left ventricular internal diameter at end systole, LVPWs = left ventricular posterior wall thickness at end systole, MI = myocardial infarction, PBS = phosphate buffered saline, rIL-4 = recombinant IL-4.

**Table 7:**
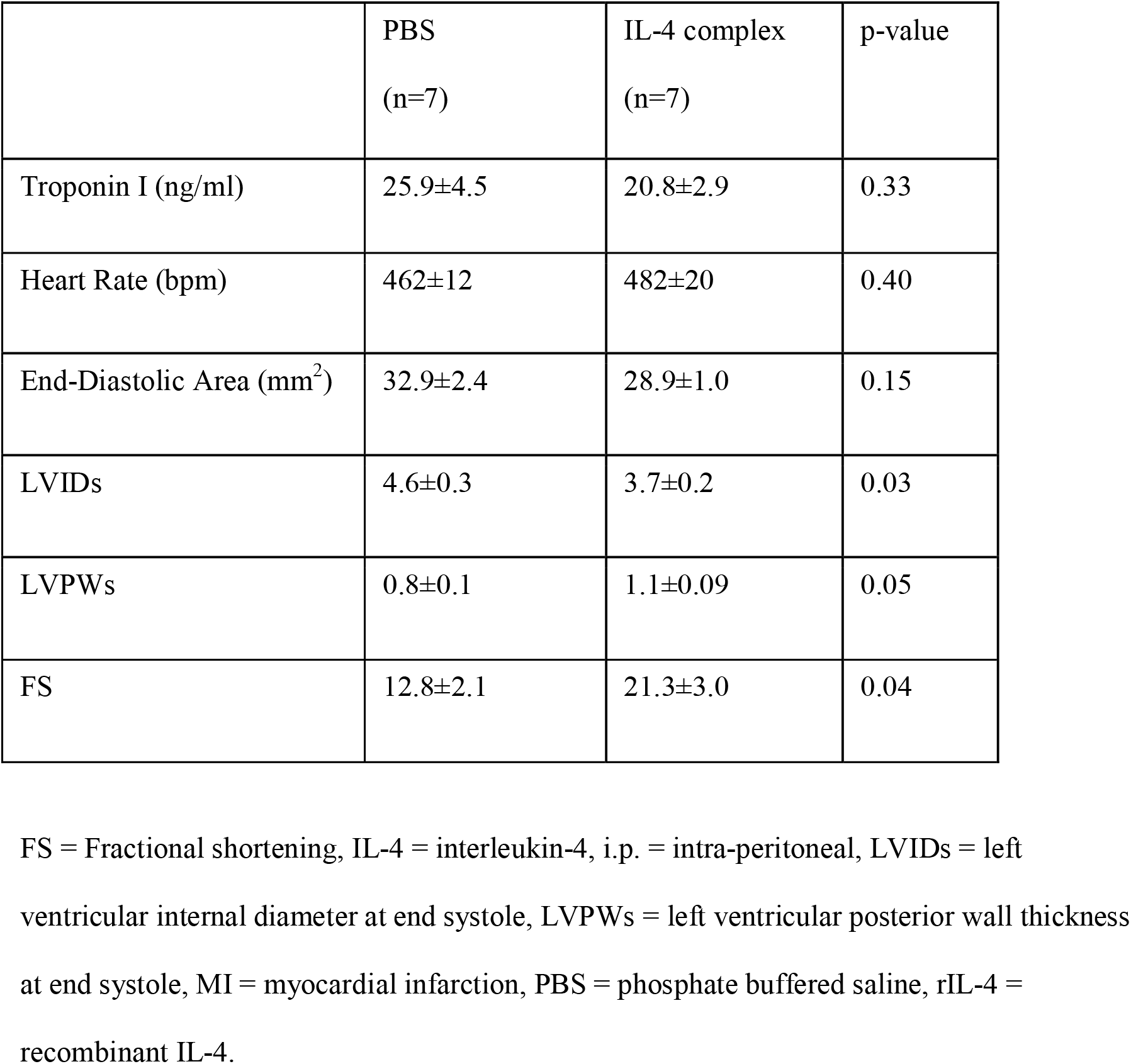
Cardiac function at Day 7 following MI in ΔdblGATA mice injected i.p. with PBS or IL-4 complex containing 5μg rIL-4 on Day 1 post-MI.

**Table 8:** Primer ID list.

| Gene | Gene name | Assay ID |
| --- | --- | --- |
| Chemokine (C-C motif) ligand 5 | Ccl5 | Mm01302427_m1 |
| Chitinase-like 3 | Chil3 | Mm00657889_mH |
| Collagen type Iα1 | Col1a1 | Mm00801666_g1 |
| Collagen type IIIα1 | Col3a1 | Mm00802300_m1 |
| Elastin | Eln | Mm00514670_m1 |
| Interleukin-1β | Il1b | Mm00434228_m1 |
| Interleukin-6 | Il6 | Mm00446190_m1 |
| Interleukin-18 | Il18 | Mm00434226_m1 |
| Interferon-γ | Ifng | Mm01168134_m1 |
| Macrophage mannose receptor | Mmr | Mm00443781_m1 |
| Matrix metalloproteinase 2 | Mmp2 | Mm00439498_m1 |
| Procollagen-lysine,2-oxoglutarate<br>5-dioxygenase 2 | Plod2 | Mm00478767_m1 |
| Resistin-like molecule $\alpha$ | Retnla | Mm00445109_m1 |
| Ribosomal protein L32 | Rpl32 | Mm02528467_g1 |
| Tissue inhibitors of matrix<br>metalloprotease 3 | Timp3 | Mm00441826_m1 |
| Transforming growth factor, beta 3 | Tgfb3 | Mm00436960_m1 |
| Tumour necrosis factor-α | Tnfa | Mm00443258_m1 |

### Tissue Cytokine Assay

Left ventricular infarct zone tissue was collected 4 days after induction of MI and snap frozen. Tissue cytokines in the heart protein extracts were assayed using the mouse Th2 Panel LEGENDplex assay (BioLegend), according to the manufactures instructions.

### Statistics

Continuous variables are expressed as mean ± standard error of mean. Continuous variables with a normal distribution were analysed using Student *t*-test, and variables with a nonnormal distribution were analysed using Wilcoxon rank sum test. Two-sided p<0.05 was considered statistically significant. All statistical analyses were performed with SPSS 21.0 statistical software package (SPSS Inc, Chicago, IL).

## RESULTS

### Eosinophils are depleted from the blood and accumulate in the myocardium of patients after acute MI

In 732 patients presenting with STEMI (Online Table 1), blood eosinophil counts declined over the first 7-9 hours after the onset of chest pain and were suppressed compared to patients with stable angina (Figure 1a, p<0.01). Peripheral blood neutrophil count in STEMI patients remained elevated at a similar level following the onset of chest pain (Figure 1a, inset), while there continued to be a decline in eosinophil count over the same time period. Histological staining of three human post-mortem hearts, collected from patients who died within 24 hours of MI (Online Table 2), revealed the presence of eosinophil peroxidase immunopositive eosinophils within the infarct area, consistent with eosinophils recruitment to the heart in humans during the acute inflammatory phase following MI (Figure 1b).

**Figure 1:**
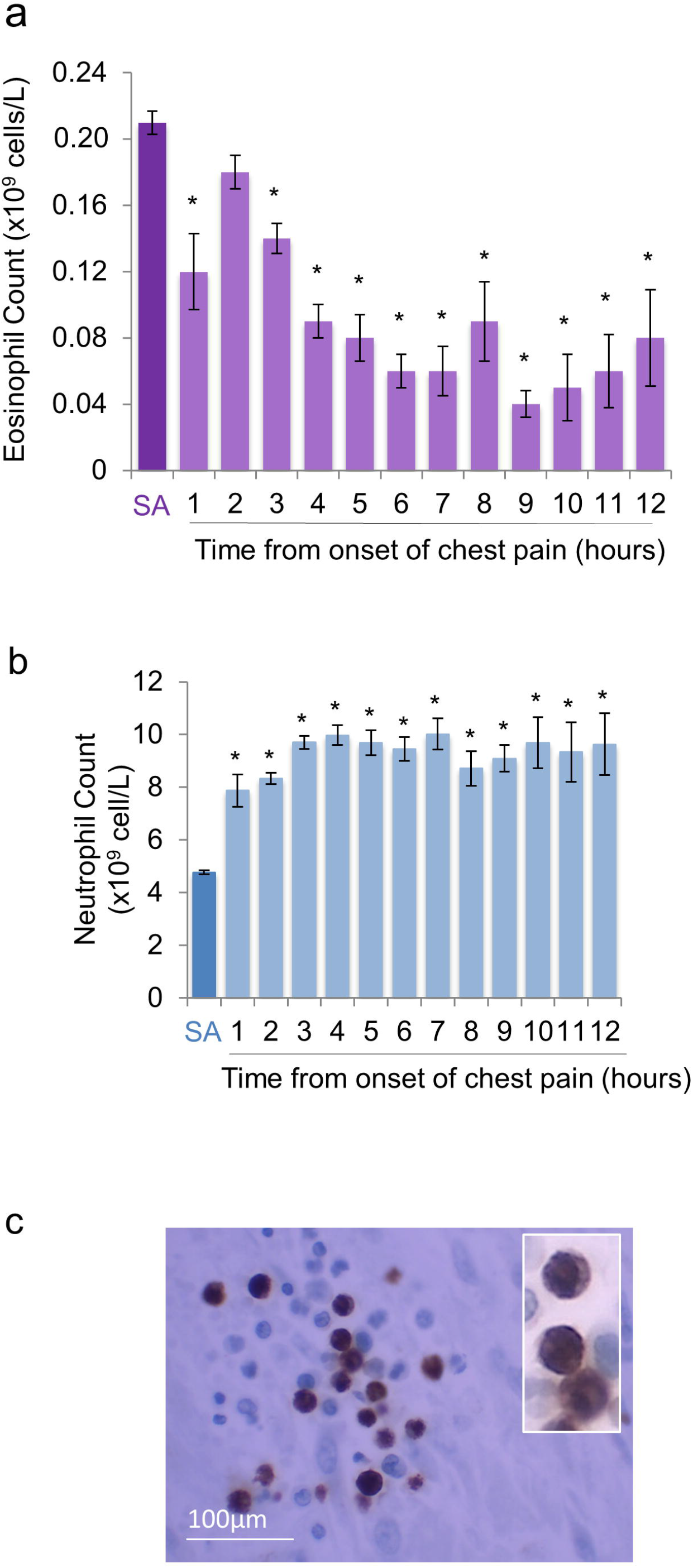
Blood eosinophil count is reduced, independently of changes in neutrophil count in STEMI patients, while eosinophils accumulate in the human heart following MI. **(a)** Peripheral blood eosinophil count and **(b)** neutrophil count in patients with stable angina (SA) or at time from onset of chest pain in patients with ST-elevation MI. *p<0.05: relative to patients with SA. Stable angina patients: n=307; ST-elevation MI patients: n=732. **(c)** Eosinophil peroxidase immunopositive eosinophils with segmented nucleus (inset) within the infarct zone of human postmortem hearts, typical of n=3.

### Eosinophils are activated following experimental MI and accumulate in the infarcted left ventricle

To permit further interrogation of the role of eosinophils in the heart following MI, eosinophil numbers in the blood and heart were also investigated in an experimental mouse model of MI following coronary artery ligation. Flow cytometric evaluation of Siglec-F+ eosinophils in blood and single cell digests of left ventricle (Figure 2a) revealed that, in parallel with clinical observations, there was a peripheral blood eosinopenia after experimental MI (p=0.003; Figure 2b). In keeping with previous studies,^12^ Siglec-F^+^Ly6G^int^ eosinophils were rarely found in uninjured hearts (Figure 2c). However, eosinophils were recruited to the heart from Day 1 post-MI, and particularly to the infarct zone, where their numbers peaked at Day 4 post-MI, during infarct repair (Figure 2c). The presense of eosinophils in tissue at day 4 was confirmed by their characteristic morphology on electron microscopy, with crystalloid containing granules in the spleen (Figure 2d) and in the infarcted heart (Figure 2e). In the infarct, variable granule morphology is consistent with eosinophil activation (Figure 2e, inset). Activation of recruited eosinophils was confirmed by flow cytometry that showed a higher intensity of Siglec-F expression on cells infiltrating the inflamed myocardium relative to naïve eosinophils residing in splenic lymphoid tissue (Figure 2f & 2g).^13^

**Figure 2:**
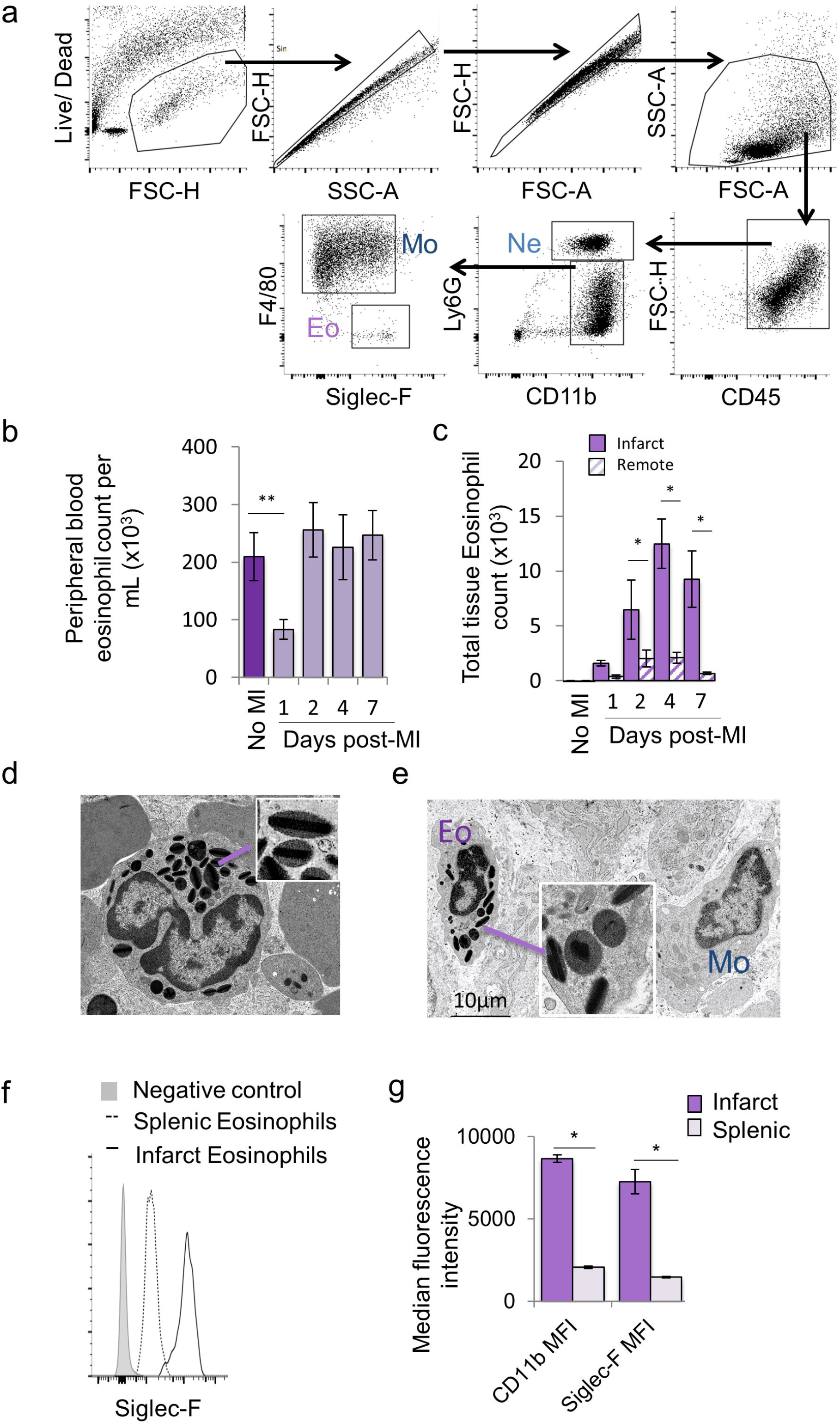
Eosinophils are reduced in the blood and accumulate in the heart following experimental MI in mice. **(a)** Representative flow cytometry plots showing the gating strategy applied to the left ventricle of WT BALB/c mice for detection of neutrophils (Ne), macrophages (Mo) and eosinophils (Eo). **(b)** Peripheral blood eosinophil count in BALB/c mice following MI (no MI: n=5; other time points: n= 5-12/group). **(c)** Total number of Siglec-F^+^Ly6G^int^ eosinophils detected by flow cytometry in the infarct and remote zones following MI (no MI: n=3; other time points: n= 5-9/group). Electron microscopy of **(d)** queiscent eosinophils in spleen, with typical electron dense crystalloid containing granules (inset) and **(e)** eosinophil (Eo) adjacent to a macrophage (Mo) in infarct, varied granule morphology is consistent with eosinophil activation (inset) **(f)** Representative histogram showing Siglec-F staining of splenic and infarct zone eosinophils at Day 4 post-MI. **(g)** CD11b and Siglec-F MFI of infarct and splenic eosinophils at Day 4 post-MI (n=4 /group). MFI = Median Fluorescence Intensity, MI = myocardial infarction, OCT = optimal cutting temperature, SA = stable angina, WT = wild type. *p<0.05, **p<0.01.

### Infarct expansion and detrimental post-MI remodelling are enhanced following genetic depletion of eosinophils

To investigate the role of eosinophils recruited to the heart following injury, MI was induced in ΔdblGATA mice with genetic deficiency of eosinophils^8^ In keeping with previous findings,^8^ analysis of peripheral blood showed no significant differences between WT BALB/c and ΔdblGATA mice with respect to white blood cell counts at baseline (Online Figure 1). Deficiency of Siglec-F^+^Ly6G^int^ eosinophils in the infarct and remote zones of the left ventricle, as well as spleen, of ΔdblGATA mice was confirmed by flow cytometry (Figure 3a). Left ventricular function and geometry were similar in ΔdblGATA and WT BALB/c mice prior to induction of MI (Online Table 3). Following the induction of MI, high-resolution ultrasound showed that hearts from ΔdblGATA mice were more dilated (Figure 2b, increased left ventricular area, p=0.02 & Online Table 4) and had greater impairment of left ventricular function than WT BALB/c mice (Figure 3c, left ventricular ejection fraction, p=0.04). Plasma troponin I concentration at 24 hours post-MI was comparable between ΔdblGATA (25.7±4.7ng/ml) and WT BALB/c mice (27.0± 3.6 ng/ml, p=0.83), indicating similar initial myocardial injury. However, by Day 7 following induction of MI, scar size was larger in ΔdblGATA mice (Figure 3d). There was no influence of eosinophil depletion on the extent of angiogenesis post-MI (Figure 3e). However, picrosirius red staining under polarizing light revealed that the proportion of tightly packed collagen fibers in the infarct zone, which have yellowish-red birefringence, was reduced in ΔdblGATA mice (Figure 3f). This was associated with an increased preponderance of thin collagen fibers, which appear green under polarized light, in the infarct zone of ΔdblGATA mice (Figure 3f). Infarct zone whole tissue qPCR at Day 7 (Figure 3g) for genes associated with post-translational collagen processing showed increased mRNA expression of procollagen-lysine, 2-oxoglutarate 5-dioxygenase 2 (*plod2*)(p<0.01), lysyl hydroxylase 2, Lysyl oxidase (*Lox*) (p<0.05) and transforming growth factor, beta 3 (*tgfb3*) (p<0.01) in ΔdblGATA mice in comparison to WT BALB/c mice. mRNA expression of Elastin (*Eln*) in the infarct zone of ΔdblGATA mice was comparable to WT BALB/c mice. These findings support a role for eosinophils in collagen scar maturation following MI.

**Figure 3:**
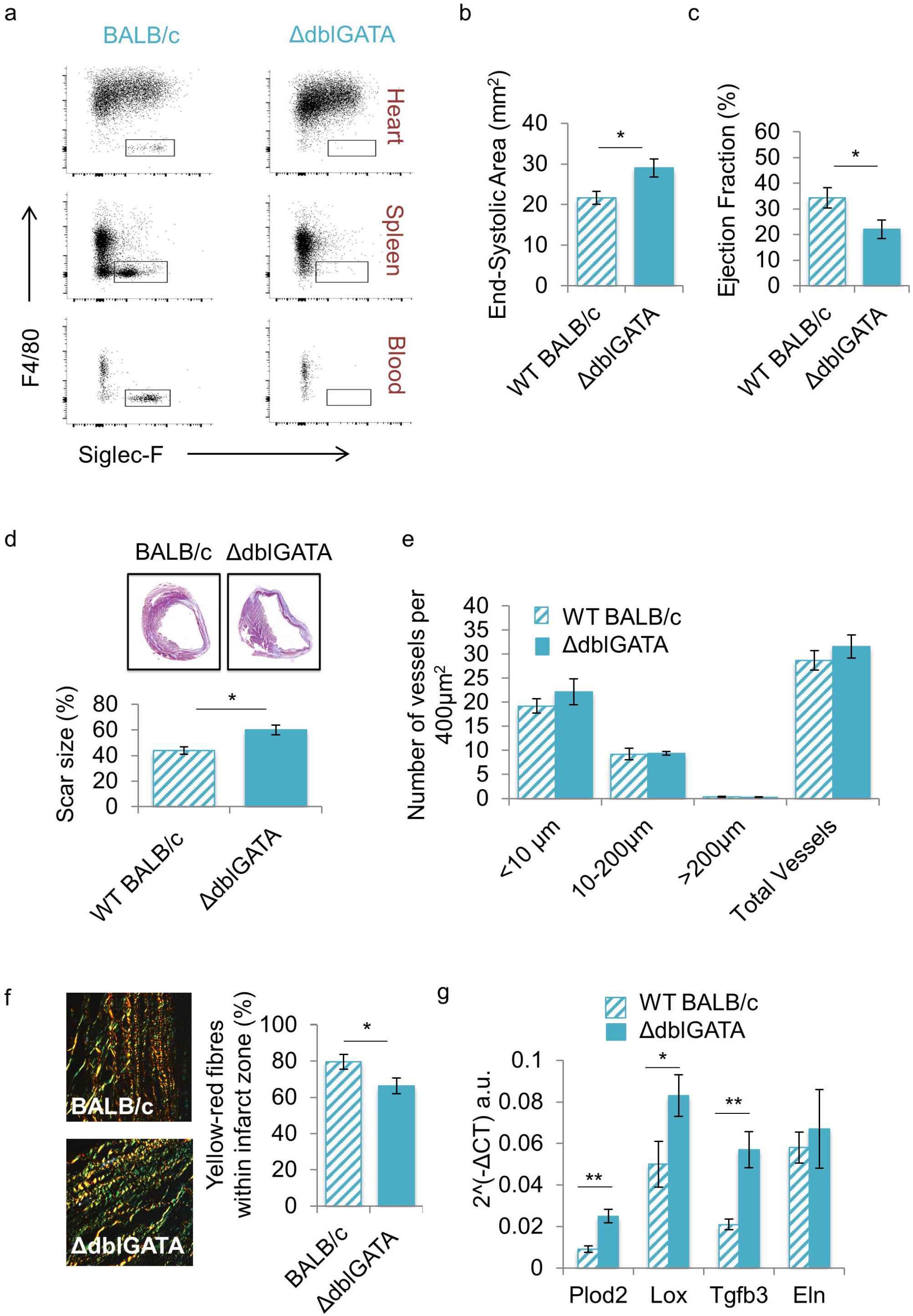
Genetic depletion of eosinophils results in enhanced detrimental remodeling post-MI accompanied by alteration of collagen scar size, morphology and collagen processing genes. **(a)** Representative flow cytometry plots demonstrating (Siglec-F^+^Ly6G^int^F4/80^-^) eosinophil deficiency in the heart, spleen and blood of ΔdblGATA mice in comparison with BALB/c mice. **(b)** End-systolic area and **(c)** Ejection fraction at Day 7 following MI in WT BALB/c and ΔdblGATA mice (n=8-9 /group). **(d)** Masson’s Trichrome stained sections for assessment of scar size, expressed in histogram as a percentage of the left ventricle. **(e)** CD31^+ve^ vessels of varying size in the infarct zone expressed per 400μm^2^ (n=6-9 /group). Imaging of picrosirius red stained collagen fibers in the infarct zone under polarised light **(f)** reveals a reduced proportion of collagen fibres in the infarct zone with yellow-red birefringence (n= 7-8/group). **(g)** mRNA expression levels of genes expressing collagen processing genes: Procollagen-Lysine, 2-Oxoglutarate 5-Dioxygenase 2 (*Plod2*) and Lysyl oxidase (*Lox*); Transforming Growth Factor beta 3 (*Tgfb3*) that regulates Plod 2 expression; and of elastin (*Eln*) in the infarct zone at Day 7 following MI (n=5 /group). *p<0.05; **p<0.01.

### Pharmacological intervention confirms enhancement of detrimental remodelling following eosinophil depletion

To confirm the data obtained with genetic deficiency of eosinophils in BALB/c mice, an antibody-mediated approach was used as an alternative means of depleting eosinophils following MI in C57BL/6 mice (Figure 4a). Successful depletion of Siglec-F^+^Ly6G^int^ eosinophils using anti-Siglec-F antiserum was confirmed by flow cytometry of single cell digests of infarcted hearts at Day 4 post-MI (Figure 4b). Depletion of eosinophils did not affect initial injury induced by MI, as measured by plasma troponin I concentrations at 24 hours post-MI (Online Table 5). High-resolution ultrasound showed that hearts from anti-Siglec-F antiserum treated mice were more dilated (Figure 4c) and had greater impairment of left ventricular function than pre-immune serum treated mice (Figure 4d, Online Table 5). Thus, pharmacological depletion of eosinophils in C57BL/6 mice confirmed the adverse cardiac remodelling phenotype associated with eosinophil depletion, further supporting a role for eosinophils in post-MI remodelling.

**Figure 4:**
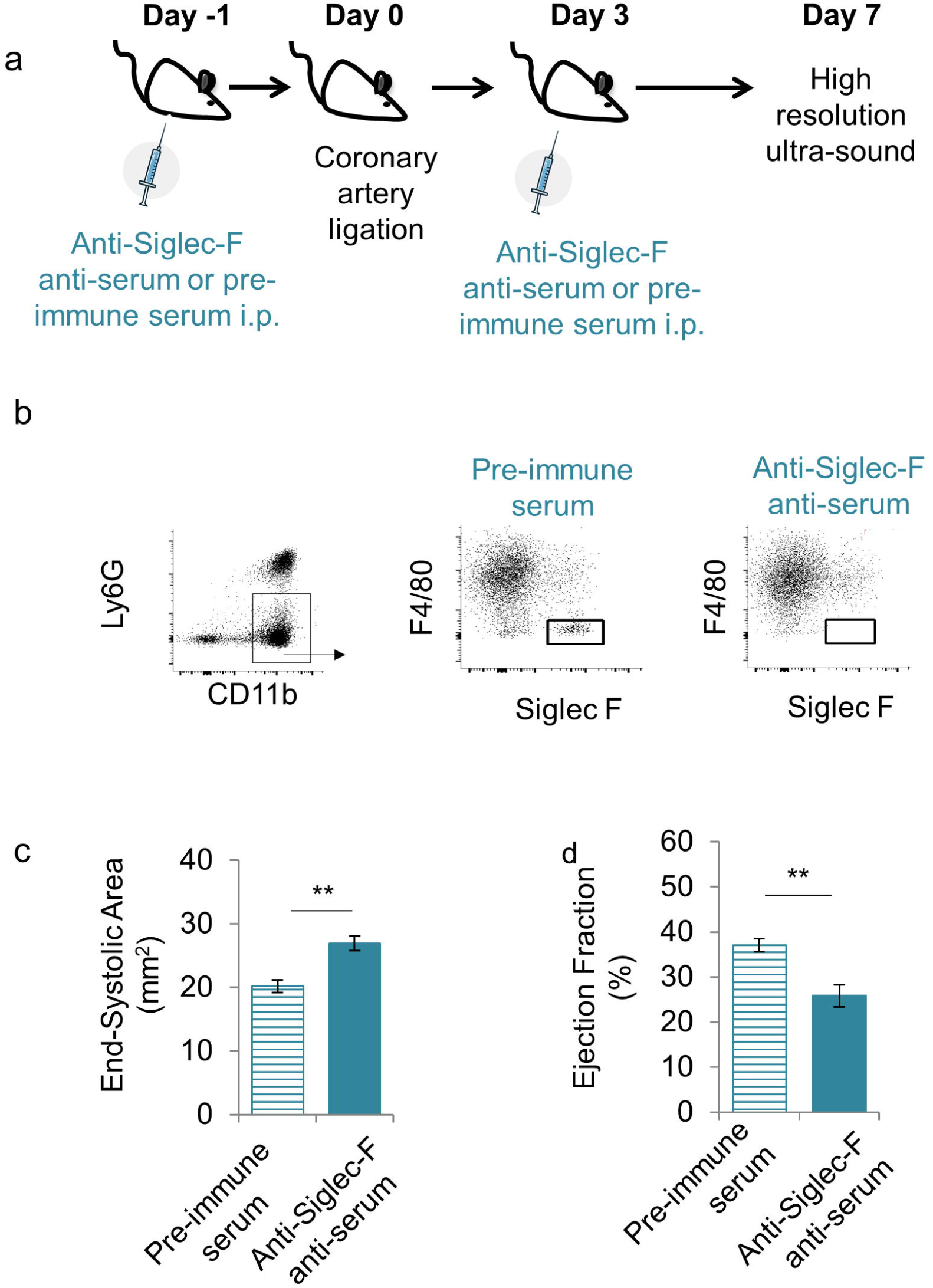
Pharmacologic depletion of eosinophils reproduces adverse remodeling in C57Bl/6 mice. **(a)** C57BL/6 mice were treated with anti-Siglec-F anti-serum, 1 day before and 3 days after induction of MI. Flow cytometry **(b)** reveals successful depletion of SiglecF+Ly6G^int^ eosinophils within the infarcted heart following antiserum treatment. **(c)** End-systolic area and **(d)** Ejection fraction at Day 7 following MI in pre-immune serum treated (n=6) and anti-Siglec-F anti-serum treated C57BL/6 mice (n=6). i.p. = intra-peritoneal. *p<0.05, **p<0.01.

### Eosinophils support an anti-inflammatory environment in the infarct zone

Anti-inflammatory Th2-type cytokines have a key role in the resolution of inflammation and are elevated in the infarct zone at Day 4 post-MI.^14^ As eosinophils are known to support a type 2 immune environment during tissue repair and regeneration we next aimed to investigate the influence of eosinophil depletion on type 2 immune mediator availability in the infarct. In keeping with an anti-inflammatory role for eosinophils ELISA showed that the infarct tissue of ΔdblGATA mice had significantly reduced availability of IL-4, IL-5, IL-13, and IL-10 (Figure 5a), in comparison to WT BALB/c mice at Day 4 following MI. In contrast, expression of genes encoding the pro-inflammatory mediators IL-18 (*Il18*), chemokine (C-C motif) ligand 5 (*Ccl5*) and tumour necrosis factor-α (*tnfa*) were all elevated in the infarct zone of ΔdblGATA mice compared to WT BALB/c mice at Day 7 post-MI (Figure 5b). Characterisation of myocardial inflammatory cell content using flow cytometry (Figure 5c) showed that the number of infarct zone Ly6G^hi^Ly6C^int^ neutrophils was higher in ΔdblGATA at Day 4 post-MI, relative to WT BALB/c mice (Figure 5d, p=0.03). Infarct zone whole tissue qPCR for genes associated with neutrophil recruitment showed increased mRNA expression of both chemokine (C-X-C motif) ligand 1 (*Cxcl1*) (p<0.05) and chemokine (C-X-C motif) ligand 2 (*Cxcl2*, p<0.05, Figure 5e). Infarct zone F4/80+ macrophage numbers were also increased in ΔdblGATA mice (Figure 5f; p=0.02) following MI. Together these data demonstrate impaired resolution of inflammation in the absence of eosinophils.

**Figure 5:**
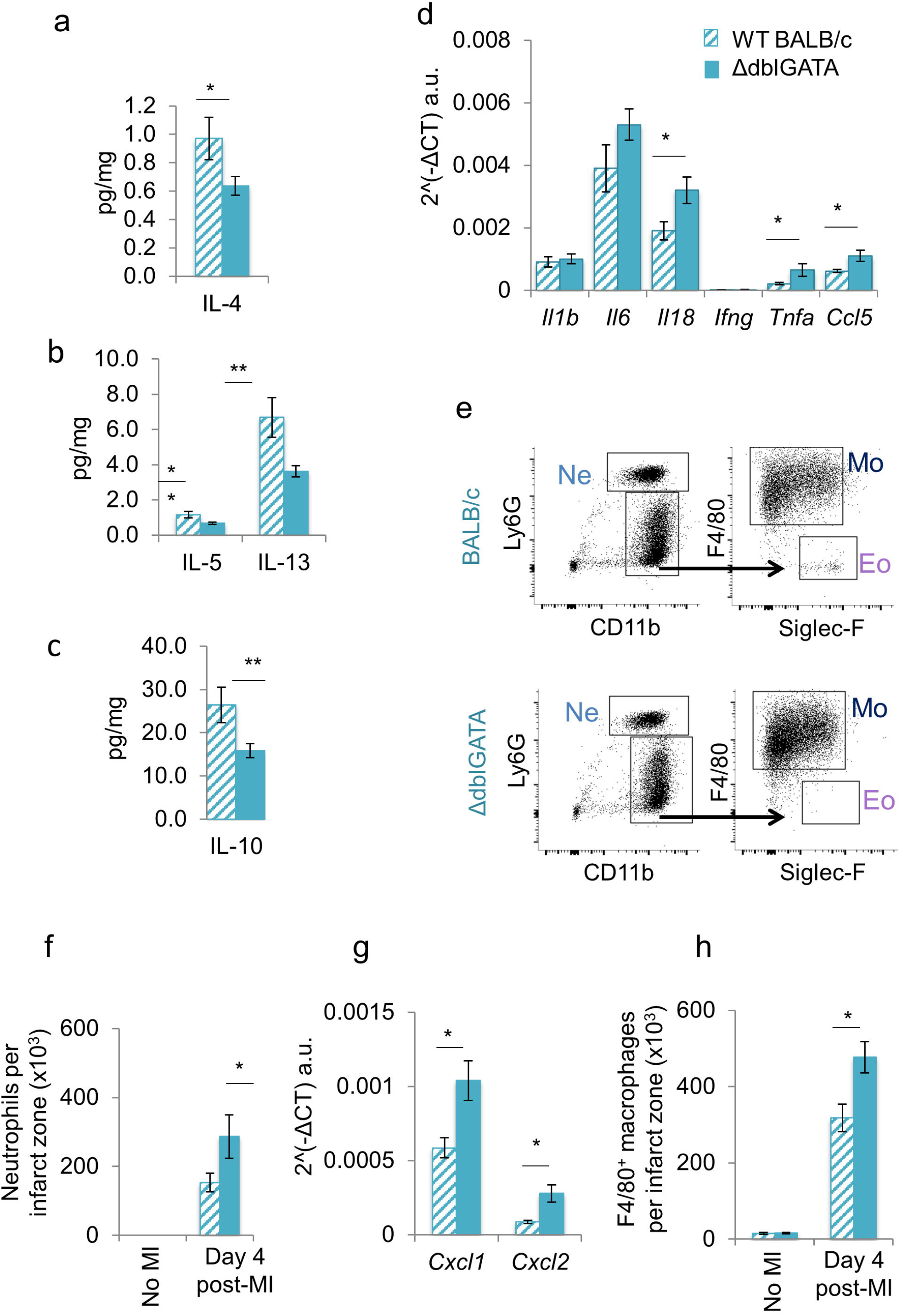
Eosinophils are required for acquisition of an anti-inflammatory Th2 dominant environment during infarct repair. ELISA reveals reduced availability of Th2 cytokines **(a)** IL-4, **(b)** IL-5 & IL-13 and **(c)** IL-10 in the infarct zones of ΔdblGATA compared to WT mice at Day 4 following MI (n=9 /group); and **(d)** increased mRNA expression of pro-inflammatory Interleukin-1β (Il1b), Interleukin-6 (Il6), Interleukin-18 (Il18), Interferon-γ (Ifng), Tumour necrosis factor-α (Tnfa), chemokine (C-C motif) ligand 5 (*Ccl5*) in the infarct zone at Day 7 following MI (n=5 /group). mRNA expression levels were normalized for Rpl32 (house keeping gene) expression. **(e)** Representative flow cytometry plots showing the gating strategy applied to the infarct zone tissue of WT BALB/c and ΔdblGATA mice for detection of neutrophils (Ne), macrophages (Mo) and eosinophils (Eo). **(f)** Total number of neutrophils in the uninfarcted heart (no-MI) and in the infarct zone 4 days after MI. (g) mRNA expression levels of Chemokine (C-X-C motif) ligand 1 (Cxcl1) and Chemokine (C-X-C motif) ligand 2 (*Cxcl2*) in the infarct zone following MI (n=5 /group). mRNA expression levels were normalized for Rpl32 (house keeping gene) expression. **(h)** Total number of F4/80+ macrophages in the uninfarcted heart (no-MI) and in the infarct zone 4 days after MI. (no MI: n=3; Day 4 post-MI: n= 7-10/group). *p<0.05, **p<0.01.

### Delayed acquisition of a pro-repair macrophage phenotype in eosinophil deficient myocardium is rescued by eosinophil replacement

Resolution of inflammation following MI occurs with a transition towards an antiinflammatory macrophage phenotype that is essential for repair and prevention of adverse cardiac remodelling.^2^ CD206 expression is typically increased on these alternatively activated anti-inflammatory macrophages.^2^ The next aim was therefore to investigate whether eosinophils were required for acquisition of a CD206^+^ anti-inflammatory macrophage phenotype following MI. CD206 was present on nearly 80% of macrophages in naïve hearts (Figure 6a). Within the first 24 hours after induction of MI, representation of CD206^+^ macrophages was reduced to less than 20% and F4/80^+^CD206^-^ pro-inflammatory macrophages dominated in the infarct zone (Figure 6a). On Day 4 post-MI, nearly 40% of infarct zone macrophages from WT BALB/c once again expressed CD206 (Figure 6a). However, acquisition of CD206 expression by infarct zone macrophages was impaired at Day 4 post-MI in ΔdblGATA mice (Figure 6a), supporting key roles for eosinophils in driving infarct zone macrophage polarisation towards an anti-inflammatory phenotype following MI. Expression of the mRNA encoding the anti-inflammatory macrophage marker, Resistin-like molecule α (*Retnla*/RELMα), was also reduced in macrophages sorted from the infarct zones of ΔdblGATA (p<0.05) compared to BALB/c mice (Figure 6b, n=4-8 /group).

**Figure 6:**
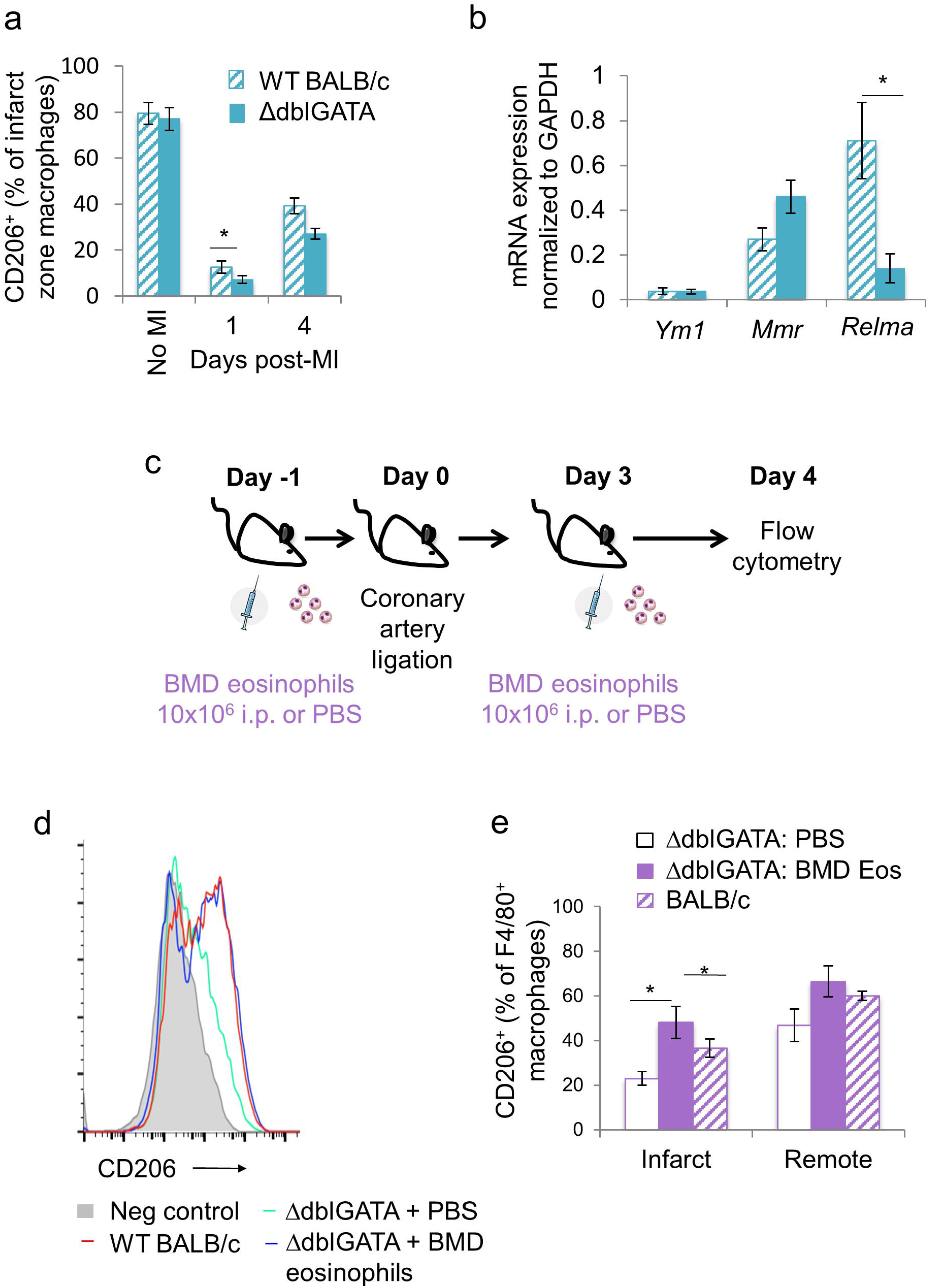
Delayed acquisition of a pro-repair macrophage phenotype in eosinophil deficient myocardium is rescued by eosinophil replacement. **(a)** The proportion of CD206 expressing F4/80^+^ macrophages in unoperated hearts (no-MI) and in the infarct zone at Day 1 & 4 post-MI in ΔdblGATA compared to WT BALB/c mice (no MI: n=3; other time points: n=4-10 /group). **(b)** mRNA expression levels of: chitinase-like 3 (*Ym1*), macrophage mannose receptor (*Mmr*) and Resistin-like molecule α (*Relma*) in macrophages sorted from the infarct zone at Day 7 following MI (n=4-8 /group). **(c)** Protocol for delivery of bone-marrow derived (BMD) eosinophils, or phosphate buffered saline (PBS), 1 day before and 3 days after MI. **(d)** Representative histogram of CD206 expressing F4/80^+^ macrophages in the infarct zones of PBS treated or BMD eosinophil replenished ΔdblGATA mice at Day 4 post-MI. **(e)** Proportion of CD206 expressing F4/80+ macrophages in the infarct and remote zones of PBS treated or BMD eosinophil replenished ΔdblGATA mice at Day 4 post-MI (n=5-7 /group). *p<0.05.

To confirm that loss of macrophage polarisation during infarct repair was due to eosinophil deficiency, bone marrow-derived (BMD) eosinophils were adoptively transferred by intra-peritoneal injection into ΔdblGATA mice before and after MI (Figure 6c). Adoptive transfer of BMD eosinophils resulted in restoration of CD206 expression on infarct zone macrophages from ΔdblGATA mice to levels of those from WT BALB/c mice (Figure 6d & 6e).

### Adverse cardiac remodelling associated with eosinophil deficiency is rescued by therapeutic IL-4 complex

As Type 2 cytokine availability is reduced in the setting of eosinophil deficiency in ΔdblGATA mice, it was reasoned that therapeutic application of IL-4 might be an effective means to rescue adverse remodelling. A long-acting IL-4 complex was therefore administered to ΔdblGATA and WT BALB/c mice 24 hours after induction of MI (Figure 7a). Plasma troponin I concentrations at 24 hours post-MI were comparable between WT BALB/c and ΔdblGATA mice prior to treatment with either PBS or IL-4 complex (Online Table 6 & 7), indicating similar initial injury in all groups. High-resolution ultrasound showed that while IL-4 complex had no impact on remodelling in eosinophil replete WT BALB/c mice when given 24 hours after injury (Figure 7b & 7c; Online Table 6), it was able to rescue the adverse remodelling phenotype in eosinophil-deficient ΔdblGATA mice (Figure 7d & 7e; Online Table 7).

**Figure 7:**
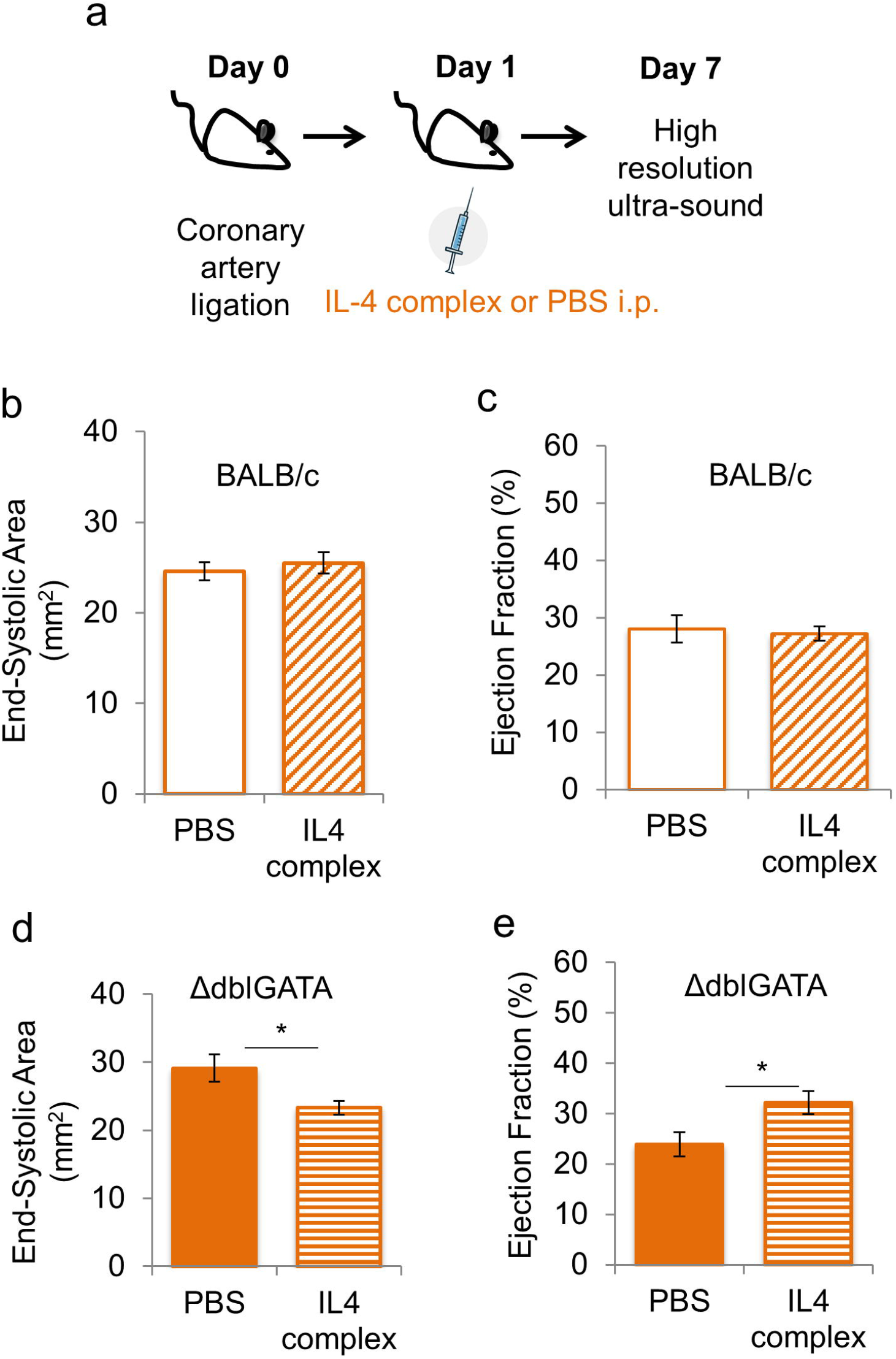
IL-4 therapy rescues the excess adverse remodelling phenotype in eosinophil deficient ΔdblGATA mice. **(a)** Protocol for administration of Interleukin-4 (IL-4) complex 1 day after induction of MI **(b)** End-systolic area and **(c)** ejection fraction at Day 7 following MI in WT BALB/c mice injected i.p. with PBS or IL-4 complex (n=5-9/group). **(d)** End-systolic area and **(h)** ejection fraction at Day 7 following MI (n=7/group) in ΔdblGATA mice injected i.p. with PBS or IL-4 complex. i.p. = intra-peritoneal, MI = myocardial infarction, PBS = phosphate buffered saline. *p<0.05; **p<0.01.

## DISCUSSION

Blood eosinophil count is reduced in the acute period following MI and we have previously shown that low peripheral blood eosinophil count predicts adverse outcome^5^. Analysis of human myocardium collected post-mortem following MI identified eosinophils, suggesting that they are recruited to the site of injury, consistent with previous identification of eosinophils in post-mortem myocardium collected following MI.^15^ However, this clinical observation leaves open the question of whether eosinophil recruitment to the injured heart indicates a negative or a positive influence on subsequent repair and remodelling. In addition it cannot be excluded that death may have influenced eosinophil recruitment to the infarct zone in these post-mortem samples. To address these questions, the role of eosinophils was investigated further using an experimental model of MI in the mouse that permits manipulation of eosinophil availability.

### Eosinophils are recruited to the heart and activated following experimental MI

In the mouse model, eosinophil numbers in the peripheral blood also declined after MI and this was associated with accumulation of eosinophils in the myocardium that began within 24 hours and continued to increase to a peak at 4 days after MI. The temporal mismatch between the decline of eosinophil numbers in the blood and accumulation in the tissue indicates a more complex regulation of peripheral blood eosinophils than can be accounted for simply by recruitment to tissue. Inflammatory cytokines produced by activated innate immune cells^16^ and stress responses, mediated by adrenal corticosteroids,^17^ can lead to eosinophil apoptosis and contribute to a decline in peripheral blood eosinophil count. Clarifying how the inflammatory environment following MI influences eosinophil survival during the period of eosinophil recruitment to the infarct zone will in future provide clues to better understand the association of peripheral blood eosinophil count with outcome in MI patients.^5–7^

Recruitment of eosinophils to the repairing mouse myocardium, reproducing the clinical observation, is consistent with an active role for eosinophils following MI. This is further supported by the observation that on becoming resident in the infarcted heart, alteration of granule morphology^4,13^ is accompaied by increased expression of Siglec-F, indicating a switch from a homeostatic to an activated phenotype.^13,18^

### Eosinophil deplation results in adverse remodelling associated with impaired scar formation

A functional role for eosinophils was confirmed *in vivo* in eosinophil-deficient ΔdblGATA mice,^8^ and also in mice with antibody-mediated eosinophil depletion with anti-Siglec-F antiserum.^13^ Both models of eosinophil depletion were accompanied by an increase in adverse structural and functional remodelling, indicating that eosinophils have a key positive impact on myocardial repair and remodelling following their recruitment from the blood. Adverse remodelling was accompanied by an increase in infarct size in the myocardium of ΔdblGATA mice relative to WT, despite no difference in the initial ischaemia-induced injury. Angiogenesis was unchanged in eosinophil deficient mice suggesting that infarct expansion did not occur due to lack of blood supply on the infarct border. Formation of a stable collagen scar is key for determination of wall stress, infarct expansion and subsequent ventricular remodelling following MI. Investigation of the infarct scar using birefringence microscopy revealed modification of collagen fibril formation in eosinophil deficient mice. Eosinophil depletion also resulted in increased infarct zone expression of *plod2*, encoding lysyl hydroxylases 2 (LH2), and also *tgfb*3, encoding Transforming Growth Factor beta 3 (TGF-β3) which upregulates *plod2* expression in fibroblasts.^19^ Increased *plod2* expression leads to increased post-translational collagen cross-linking, resulting in reduced tensile strength.^20–22^ The detrimental effects of increased collagen cross-linking have been identified in the weakened aortic wall present in Marfan syndrome and abdominal aortic aneurysms.^22^ Disrupted post-translational collagen processing in the infarct zone may therefore underlie infarct expansion and ventricular dilatation observed following MI in the absence of eosinophils.

### Eosinophils are required to promote resolution of inflammation during post-infarct repair

Inflammation is essential for infarct repair; but when excessive or prolonged is associated with adverse cardiac remodelling.^23^ In other settings eosinophil-derived pro-resolving lipid mediators can reduce the numbers of neutrophils in inflamed tissue,^3,4^ by counter-regulating neutrophil influx and stimulating macrophage phagocytosis of apoptotic neutrophils,^5^ thus promoting the resolution of inflammation. Neutrophil recruitment to the heart following MI was increased in eosinophil-deficient ΔdblGATA mice, and this was associated with increased expression of pro-inflammatory mediators, including neutrophil recruiting chemokines. Eosinophil-deficient ΔdblGATA mice also had reduced availability of the antiinflammatory Th2-type cytokines, IL-4, IL-5, IL-13, and IL-10, in the infarct tissue. This was accompanied by a reduction in the proportion of infarct zone macrophages expressing CD206+ and RELM-α, that identify anti-inflammatory, pro-resolution and pro-repair macrophage phenotype in the infarct zone.^2^ Following MI, CD206^+^ macrophages produce paracrine mediators that activate cardiac fibroblasts promoting scar formation.^2^ Consistent with our data Qin *et al*.^24^ have recently demonstrated increased expression of CD206 and RELM-α when cultured macrophages were treated with human eosinophil-conditioned media. In the current study loss of CD206^+^ macrophages in the infarct of eosinophil-deficient ΔdblGATA mice was rescued by eosinophil replenishment. Collectively, these findings support a role for eosinophils in driving transition to resolution of inflammation and stable scar formation during infarct repair.

### Therapeutic IL-4 rescues adverse post-MI in the setting of eosinophil deficiency

IL-4 mediated activation of macrophages is known to induce characteristic expression of specific effector molecules (e.g. RELM-α) and is associated with the adoption of a wound healing phenotype.^25^ Previous experiments have demonstrated the importance of this phenotype during post-MI repair^2^ and the ability of therapeutic IL-4 to enhance accumulation of CD206+ ‘wound healing’ macrophages and prevent detrimental remodelling, at least when administered immediately after injury. Eosinophils are a key source of IL-4 in a number of physiological processes, including non-cardiac tissue repair and regeneration.^18,26,27^ Experiments in eosinophil-deficient mice showed IL-4 availability in the infarct was significantly reduced at Day 4 following MI, the time point at which eosinophil recruitment to the myocardium peaks, and also a key time-point for transition of macrophages from an inflammatory to a repair phenotype.^28^ We therefore investigated the potential for IL-4 to rescue detrimental post-MI remodelling in eosinophil-deficient mice. Interestingly, administration of IL-4 complex 24 hours after MI, when eosinophil recruitment to the myocardium is already significantly increased, had no influence on post-MI remodelling in eosinophil-replete mice. This is consistent with the previous studies that showed therapeutic benefit of IL-4 only when given prior to,^2^ or immediately after MI.^3^ However, although IL-4 is only one of several mediators released by eosinophils with the potential to modify infarct repair, IL-4 complex administration was effective in reversing adverse cardiac remodelling observed in eosinophil-deficient mice.

In summary, this study identifies eosinophils as key effector cells of the type 2 innate immune response required for post-MI cardiac repair and prevention of adverse remodelling. IL-4 therapy, proposed as a means to improve outcomes post-MI,^3^ was effective in reversing detrimental outcomes in the setting of eosinophil deficiency in experimental MI. The use of biomarkers such as a persistently low peripheral blood eosinophil count post-MI may therefore provide a means to direct IL-4 therapy to patients who might gain the most benefit from it.

### Competency in medical knowledge

The resolution of inflammation is one of the central features of successful infarct healing following myocardial infarction. This process is modulated by a number of cellular and molecular mediators including eosinophils, likely through the provision of anti-inflammatory mediators, like IL-4.

### Translational Outlook

Further studies are needed to evaluate prospectively the prognostic predictive value of a low peripheral blood eosinophil count and the potential therapeutic efficacy of IL-4 in preventing adverse cardiac remodelling in patients following myocardial infarction.

**Table 9:** Antibodies for Flow cytometry.

| Monoclonal rat<br>anti-mouse ab | Clone | Fluorophores | Manufacture | Concentration |
| --- | --- | --- | --- | --- |
| CD45.2 | 104 | PE Cy7 | Biolegend | 1:100 |
| CD45.2 | 104 | BV650 | Biolegend | 1:500 |
| CD11b | M1/70 | AF700 | Biolegend | 1:200 |
| F4/80 | BM8 | PE Cy7 | BD<br>Biosciences | 1:200 |
| Ly6G | 1A8 | Pacific Blue | Biolegend | 1:200 |
| Siglec-F | E50-2440 | AF647 | BD<br>Pharminen | 1:100 |
| Siglec-F | E50-2440 | PE | BD<br>Pharminen | 1:100 |
| CD206 | C068C2 | FITC | Biolegend | 1:100 |
| Ly6C | HK1-4 | PerCP Cy5.5 | BD<br>Pharminen | 1:100 |
| CD115 | T38-320 | APC | BD<br>Pharminen | 1:100 |
|  |  | DAPI | Life<br>Technologies | 1:1000 |
| anti-CD16/32 | 2.4G2 |  | BD Bioscience | 1:200 |

## Acknowledgments

We thank Chris-Anne Mackenzie and Tracey Millar at the Edinburgh Brain & Tissue Bank for help in accessing human tissue samples; Melanie McMillan, Mike Millar & Moira Lamont for support with histology & immunostaining. Shonna Johnston and Will Ramsay from the QMRI Flow Cytometry & Cell Sorting Facility and Steve Mitchell from the KB Electron Microscopy Facility for their advice & technical assistance. Professor Paul Crocker, University of Dundee, gifted both sheep anti-Siglec-F polyclonal antiserum and sheep pre-immune serum. The eosinophil peroxidase antibody was supplied by Professor Elizabeth Jacobsen, the Mayo Clinic, Scottsdale, Arizona.

## List of abbreviations

BMD: bone marrow-derived
ELISA: Enzyme linked immunoabsorbent assay
hi: high
IL: interleukin
int: intermediate
ip: intra-peritoneal
MI: myocardial infarction
lo: low
STEMI: ST-segment elevation myocardial infarction
WT: wild type

---

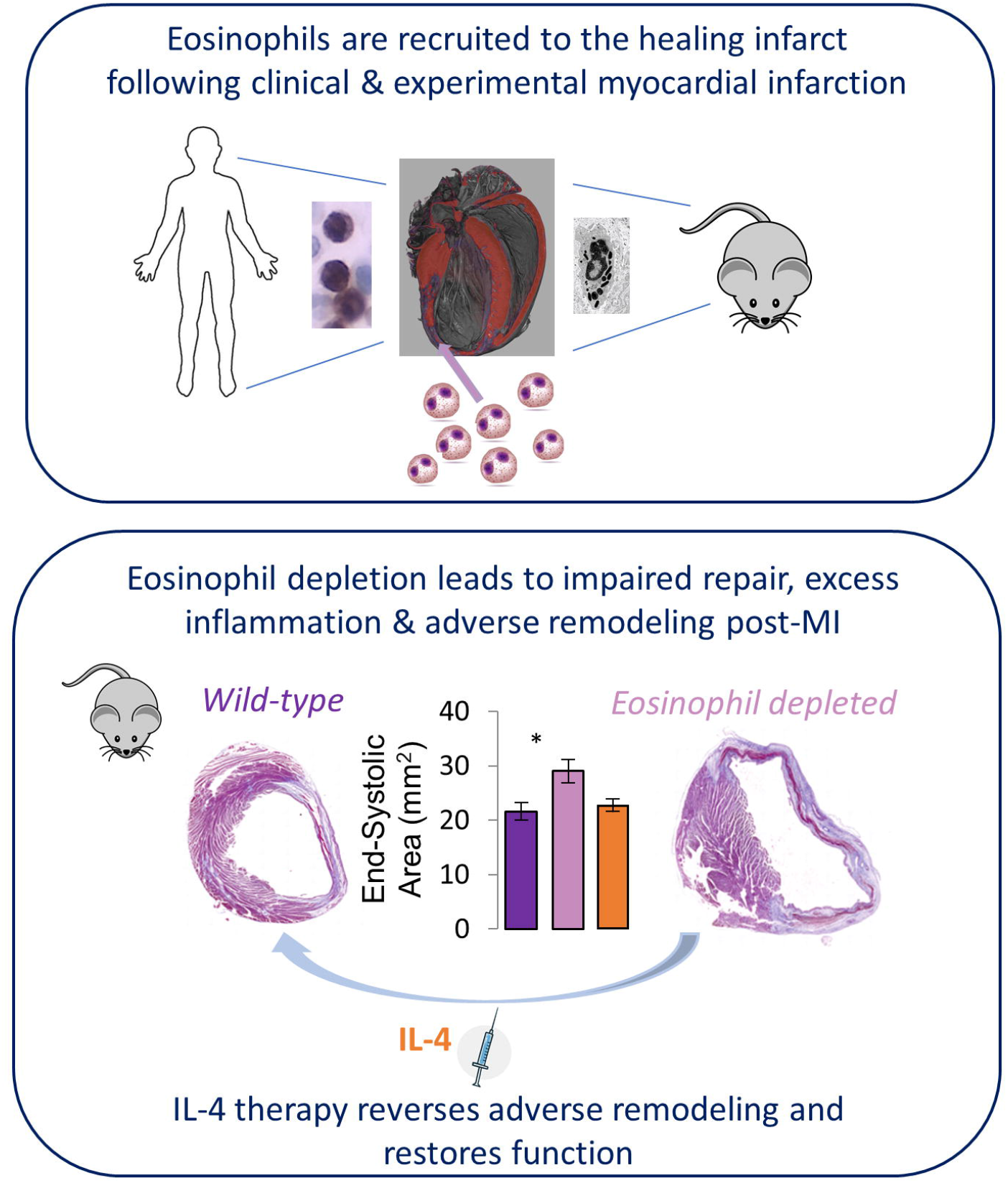

## References

1. Nahrendorf M, Swirski FK, Aikawa E, et al. The healing myocardium sequentially mobilizes two monocyte subsets with divergent and complementary functions. J Exp Med. 2007;204:3037–3047.

2. Shiraishi M, Shintani Y, Shintani Y, et al. Alternatively activated macrophages determine repair of the infarcted adult murine heart. J Clin Invest. 2016;126:2151–2166.

3. Shintani Y, Ito T, Fields L, et al. IL-4 as a Repurposed Biological Drug for Myocardial Infarction through Augmentation of Reparative Cardiac Macrophages: Proof-of-Concept Data in Mice. Sci Rep. 2017;7:6877.

4. Lee JJ, Jacobsen EA, Ochkur SI, et al. Human versus mouse eosinophils: “that which we call an eosinophil, by any other name would stain as red.” J Allergy Clin Immunol. 2012;130:572–584.

5. Toor IS, Jaumdally R, Lip GYH, Millane T, Varma C. Eosinophil count predicts mortality following percutaneous coronary intervention. Thromb Res. 2012;130:607–611.

6. Shiyovich A, Gilutz H, Plakht Y. White Blood Cell Subtypes Are Associated with a Greater Long-Term Risk of Death after Acute Myocardial Infarction. Tex Heart Inst J. 2017;44:176–188.

7. Konishi T, Funayama N, Yamamoto T, et al. Prognostic Value of Eosinophil to Leukocyte Ratio in Patients with ST-Elevation Myocardial Infarction Undergoing Primary Percutaneous Coronary Intervention. J Atheroscler Thromb. 2017;24:827–840.

8. Yu C, Cantor AB, Yang H, et al. Targeted deletion of a high-affinity GATA-binding site in the GATA-1 promoter leads to selective loss of the eosinophil lineage in vivo. J Exp Med. 2002;195:1387–1395.

9. Thygesen K, Alpert JS, Jaffe AS, et al. Third universal definition of myocardial infarction. Eur Heart J. 2012;33:2551–2567.

10. Mohrs M, Ledermann B, Köhler G, Dorfmüller A, Gessner A, Brombacher F. Differences Between IL-4- and IL-4 Receptor α-Deficient Mice in Chronic Leishmaniasis Reveal a Protective Role for IL-13 Receptor Signaling. J Immunol. 1999;162:7302–7308.

11. White CI, Jansen MA, McGregor K, et al. Cardiomyocyte and vascular smooth muscle-independent 11β-hydroxysteroid dehydrogenase 1 amplifies infarct expansion, hypertrophy, and the development of heart failure after myocardial infarction in male mice. Endocrinology. 2016;157:346–357.

12. Diny NL, Diny NL, Hou X, et al. Macrophages and cardiac fibroblasts are the main producers of eotaxins and regulate eosinophil trafficking to the heart. Eur J Immunol. 2016;46:2749–2760.

13. Griseri T, Arnold IC, Pearson C, et al. Granulocyte Macrophage Colony-Stimulating Factor-Activated Eosinophils Promote Interleukin-23 Driven Chronic Colitis. Immunity. 2015;43:187–200.

14. Timmers L, Sluijter JPG, Van Keulen JK, et al. Toll-like receptor 4 mediates maladaptive left ventricular remodeling and impairs cardiac function after myocardial infarction. Circ Res. 2008;102:257–264.

15. Atkinson JB, Robinowitz M, McAllister HA, Virmani R. Association of eosinophils with cardiac rupture. Hum Pathol. 1985;16:562–568.

16. Bass DA. Behavior of eosinophil leukocytes in acute inflammation. II. Eosinophil dynamics during acute inflammation. J Clin Invest. 1975;56:870–879.

17. Bass DA, Gonwa TA, Szejda P, Cousart MS, DeChatelet LR, McCall CE. Eosinopenia of acute infection: Production of eosinopenia by chemotactic factors of acute inflammation. J Clinical Invest. 1980;65:1265–1271.

18. Wu D, Molofsky AB, Liang H-E, et al. Eosinophils Sustain Adipose Alternatively Activated Macrophages Associated with Glucose Homeostasis. Science. 2011; 332:243–247.

19. Van Der Slot AJ, Van Dura EA, De Wit EC, et al. Elevated formation of pyridinoline cross-links by profibrotic cytokines is associated with enhanced lysyl hydroxylase 2b levels. Biochim Biophys Acta Mol Basis Dis. 2005;1741:95–102.

20. Brinckmann J, Notbohm H, Tronnier M, et al. Overhydroxylation of lysyl residues is the initial step for altered collagen cross-links and fibril architecture in fibrotic skin. J Invest Dermatol. 1999;113:617–621.

21. Pornprasertsuk S, Duarte WR, Mochida Y, Yamauchi M. Overexpression of lysyl hydroxylase-2b leads to defective collagen fibrillogenesis and matrix mineralization. J Bone Miner Res. 2005;20:81–87.

22. Ploeg M, Gröne A, van de Lest CHA, et al. Differences in extracellular matrix proteins between Friesian horses with aortic rupture, unaffected Friesians and Warmblood horses. Equine Vet J. 2016;1–5.

23. Panizzi P, Swirski FK, Figueiredo J, et al. Impaired infarct healing in atherosclerotic mice with Ly-6Chi monocytosis. J Am Coll Cardiol. 2010;55:1629–1638.

24. Qin M, Wang L, Li F, et al. Oxidized LDL activated eosinophil polarize macrophage phenotype from M2 to M1 through activation of CD36 scavenger receptor. Atherosclerosis. 2017;263:82–91.

25. Ruckerl D, Allen JE. Macrophage proliferation, provenance, and plasticity in macroparasite infection. Immunol Rev. 2014;262:113–133.

26. Goh YPS, Henderson NC, Heredia JE, et al. Eosinophils secrete IL-4 to facilitate liver regeneration. Proc Natl Acad Sci USA. 2013;110:9914–9919.

27. Heredia JE, Mukundan L, Chen FM, et al. Type 2 innate signals stimulate fibro/adipogenic progenitors to facilitate muscle regeneration. Cell. 2013;153:376–388.

28. Hilgendorf I, Gerhardt LMS, Tan TC, et al. Ly-6 chigh monocytes depend on nr4a1 to balance both inflammatory and reparative phases in the infarcted myocardium. Circ Res. 2014;114:1611–1622.

