## Supplementary Methods for "Eosinophil deficiency promotes aberrant repair and adverse remodelling following acute myocardial infarction"

**Patient selection with ST-segment elevation MI**

Exclusion criteria for STEMI patients were cardiogenic shock on admission (systolic BP of <100 mmHg and signs of acute circulatory failure), absence of a completely occluded culprit vessel (TIMI 0 flow) on admission, failure to achieve TIMI III flow following percutaneous coronary intervention, and no admission white cell differential count. Patient clinical characteristics were derived from the local Cardiac Services Database System (Phillips CVIS).

**Myocardial Infarction**

The permanent coronary artery ligation model was used to induce myocardial infarction by ligating the left anterior descending coronary artery with a suture. Mice were anaesthetized with isoflurane inhalation. Following induction of anaesthesia, buprenorphine (0.05 mg/kg) was given subcutaneously for perioperative analgesia, in addition to 1 mL 0.9% saline for rehydration. The mice were connected to a mechanical ventilator (120 breaths/minute, 240 μL stroke volume, Hugo Sachs Elektronik Minivent Harvard Apparatus). A left-sided thoracotomy performed to open the 3rd intercostal space. The left anterior descending artery visible at the left atrial notch and its superficial course over the anterior surface of the heart was identified. The pericardium was left intact. A 8.0 Ethalon (Ethicon, Livingstone) was used to ligate the left anterior descending artery in its mid course. The chest was then closed in layers with 5/0 monofilament sutures and pneumothorax evacuated. Following the return of spontaneous respiratory effort and self-righting of mice to the prone position, the mice were extubated. Animals were returned to their cage which was placed on a heat mat maintained at 37°C for 24 hours. Further buprenorphine (0.05 mg/Kg) and/or 0.9% saline was administrated the following day if necessary. For the eosinophil depletion experiments, mice were administered either sheep polyclonal antiserum raised to Siglec-F or sheep pre-immune serum (both gifted by Professor Paul Crocker, University of Dundee) via intra-peritoneal injection on Day (-1) and Day 3 following MI. Each dose of 100 μL anti-siglec-F antiserum was diluted with 100 μL sterile PBS^-/-^. The sheep preimmune serum was first sterile-filtered and each dose of 100 μL sheep pre-immune serum diluted with 100 μL sterile PBS^-/-^. For IL-4 replenishment experiments, ΔdblGATA mice were administered 5μg of recombinant murine IL-4 (Peprotech) complexed to 25 μg anti-IL-4 antibody (clone 11B11; BioXcell) in 100 μL sterile PBS without Ca2+ and Mg2+ (PBS^-/-^) or 100 μL sterile PBS^-/-^ via intra-peritoneal injection at Day 1 post-MI.

**High-resolution ultrasound measurements**

Left ventricular structure and function was assessed following MI using high-resolution ultrasound (VisualSonics Vevo 770, Toronto, Canada) with a 707B 30MHz ultrasound probe. Mice were maintained under light anaesthesia with inhaled isoflurane, ensuring that the heart rate was maintained >500 beats/min. Two-dimensional images were obtained of parasternal long-axis views of the left ventricle. The investigator was blind to group assignment. Images were saved and analyzed offline using Vevo 770 high-resolution imaging system, version 2.2.0 to calculate the left ventricular end-diastolic area (LVEDA), left ventricular end-systolic area (LVESA) and ejection fraction (EF). The image analysis was performed in a single-blinded manner.

**Cardiac Troponin I ELISA**

Twenty-fours hours after performing coronary artery ligation, tail blood samples were collected in 3.2% sodium citrate buffer and the plasma layer removed after centrifugation at 6000g for 8 minutes. Cardiac troponin I was measured in plasma samples by ELISA (Life Diagnostics Inc. High Sensitivity Mouse Cardiac Troponin I ELISA kit) to assess the size of myocardial injury. A plate reader was used to determine the concentration of cardiac troponin I which is proportional to the absorbance at 405 nm. The assays detection limit for troponin I was 0.156 ng/mL. Only mice with a plasma troponin I level greater than 10 ng/mL were regarded as having suffered a significant MI.

**Bone marrow-derived eosinophil cell culture**

Base media was prepared which contained RPMI 1640 and 20% Fetal Calf Serum (Gibco), 100 IU/mL penicillin and 10 μg/mL streptomycin (Cellgro), 25 mM HEPES, 2 mM L-glutamine, 1x non-essential amino acids and 1 mM sodium pyruvate (Life Technology), and 50μM 2-betamercaptoethanol (Sigma). Bone marrow cells were collected from the femurs and tibiae of naïve wild-type BALB/c mice by flushing the opened bones with RPMI 1640. After centrifugation, the red blood cells were lysed with sterile dH2O and then 10x PBS^-/-^ added to make the solution isotonic. Following further centrifugation, the cells were washed in PBS^- /-^. The bone marrow cells were resuspended at 1x10^6^ cells/mL in the base media supplemented with 100 ng/mL stem cell factor (SCF; PeproTech) and 100 ng/ml FLT3 ligand (FLT3-L; PeproTech) and incubated in a humidified, 5% CO_2_ atmosphere at 37°C in Nunc™ Cell Culture Treated EasYFlasks™. Every other day, from days 0 to 4, one-half of the base media was replaced with fresh base media containing SCF and FLT3-L to achieve a concentration of 1x10^6^ cells/ml. At Day-4 the base media containing SCF and FLT3-L was replaced with base media containing 10 ng/ml recombinant mouse IL-5 (rmIL-5; R&D Systems). Every other day from this point onwards, one-half of the base media was replaced with fresh base media containing rmIL-5 to achieve a concentration of 1x10^6^ cells/mL. At Day-8, non-adherent cells were transferred to a fresh Nunc™ Cell Culture Treated EasYFlasks™. On Day-0 and the days indicated, cell counts were determined using a NC-100™ Nucleocounter and 50,000 cells were taken to characterize the cells by performing a cytospin and flow cytometric analysis. The cytocentrifuge preparations were fixed and stained with Eosin and Haematoxylin. Purity of greater than 90% for eosinophils was confirmed prior to intra-peritoneal injection into ΔdblGATA mice by flow cytometric analysis. This was performed on an LSR II instrument (BD Biosciences) and data analyzed using FlowJo software (Tree Star) to calculate eosinophil purity.

**Histopathological analysis**

Samples of paraffin embedded infarcted human myocardium were obtained from the Edinburgh Brain & Tissue Bank (Online Table 2). Samples were deparaffinised in xylene then rehydrated through graded alcohols. Antigen retrieval was by pressure cooking in citric acid (0.1%). Slides were sequentially blocked with peroxidase and protein blocks prior to incubation with eosinophil peroxidase antibody (supplied by Elizabeth Jacobsen, Mayo Clinic, Scottsdale, Arizona), at 1:500 dilution. Following washing in TBS, slides were incubated with Novolink^TM^ polymer (Leica Biosystems), and peroxidase activity was developed with DAB working solution. Sections were lightly counterstained with haematoxylin & eosin, cleared and cover slipped.

Animals were euthanized and perfused at 100mmHg with heparinised saline followed by 4% neutral buffered formaldehyde. Hearts were harvested and fixed with 4% neutral buffered formaldehyde at atmospheric pressure for 18 hours and then transferred to 70% ethanol for 3 days. Following which the hearts were dissected into two parts along the short-axis at the level of the ligation suture and embedded in paraffin. From the paraffin blocks, 5 μm sections were cut using a microtome (Leica, Germany), at three different levels below the level of coronary artery ligation, each 300 μm apart.

Scar size was assessed using Masson trichrome staining, by expressing the epicardial length of the infarct as a percentage of the LV epicardial circumference. Picrosirius red staining was examined under polarized light and bright field microscopy to assess myocardial fibrosis in the infarct zone. Thick collagen fibers appeared red/yellow and thin collagen fibers appeared green under polarized light. The red/yellow fibers in each field of view were expressed as a proportion of the total collagen fibers visualized under polarized light. The images for picrosirius red and Masson trichrome staining were acquired at 20x magnification with an Axioscan slider scanner (Carl Zeiss, Germany). Subsequently, image analysis was performed using Image-Pro Plus (Image- Pro Plus, Version 9.1, Silver Spring, MD).

**Transmission Electron Microscopy**

Mice were anaesthetised and perfused at 100mmHg with heparinised saline followed by 3% glutaraldehyde in 0.1M Sodium Cacodylate buffer, pH 7.3. Heart and spleen were placed in fixative for a further 3h at room temperature prior to dissection ofsections for analysis. Specimens were then post-fixed in 1% Osmium Tetroxide in 0.1M Sodium Cacodylate for 45 minutes, then washed in three 10 minute changes of 0.1M Sodium Cacodylate buffer. Samples were dehydrated in 50%, 70%, 90% and 100% ethanol (X3) for 15 minutes each, then in two 10-minute changes in Propylene Oxide. Samples were then embedded in TAAB 812 resin. Sections, 1μm thick were cut on a Leica Ultracut ultramicrotome, stained with Toluidine Blue, and viewed in a light microscope to select suitable areas for investigation. Ultrathin sections, 60nm thick were cut from selected areas, stained in Uranyl Acetate and Lead Citrate then viewed in a JEOL JEM-1400 Plus TEM. Representative images were collected on a GATAN OneView camera. Eosinophils were identified by their typical morphology with crystalloid granules.

**RNA extraction and real-time quantitative PCR**

RNA was extracted from infarct zone whole tissue as per manufacturer’s instructions using the RNeasy® Mini Kit, Part 1 (January 2011, QIAGEN) protocol. RNA extracted from infarct zone whole tissue was reverse transcribed to single stranded cDNA using the QuantiTect Reverse Transcription Kit (QIAGEN). Reverse transcription was carried out under the following conditions: 42°C for 30 minutes and 95°C for 5 minutes. TAQman® gene expression assays (Online Table 10) were used for qRT-PCR. mRNA expression levels were normalized for Rpl32 (house keeping gene) expression and presented as fold changes. The samples were run in duplicate and averaged. If there was greater than 10% discrepancies between values then the samples were removed from the analysis.

**Cells**

30 μL tail blood samples (in 30 μL 3.2% citrate buffer) were stained with a mixture of antibodies (Online Table 10) at 4°C for 30 minutes. Subsequently, tail blood samples underwent red blood cell lysis with FACS lysing buffer (BD Bioscience, 1:10 dilution in dH2O) and were then washed PBS^-/-^. Total white blood cell numbers were determined using Flow-Check™ fluorospheres (Life Technology). Peritoneal cells were harvested by performing a peritoneal cavity lavage and triturated through a 40 μm nylon mesh (Fisherbrand). Spleens were collected from naïve mice and following MI, and prepared as a single-cell suspension by mechanical dissociation and triturated through a 40 μm nylon mesh. For both the peritoneal cells and splenocyte, the cell suspensions were centrifuged, treated with red blood cell lysing buffer (Sigma) and washed with PBS^-/-^ and total leukocyte numbers were determined. Hearts were perfused with heparinized saline and the left ventricles harvested from naïve mice and following MI. The infarcted left ventricle was dissected in order to separate it into infarct and remote zone myocardium. The tissues were transferred to gentle MACS™ tubes containing Hanks’ balanced salt solution (HBSS) with Ca^2+^ and Mg^2+^ in which was dissolved collagenase D (1.25 mg/mL) (Roche, Lakewood, NJ) and DNase I (60U U/mL) (Sigma, St. Louis, MO). Infarct and remote zone myocardium samples were homogenized using Heart protocol 1 on a MACS™ Dissociator. After dissociation, single-cell suspensions were prepared by incubating the enzyme mix at 37°C for 30 minutes whilst being gently agitated. Tissues underwent another round of mechanical dissociation using Heart protocol 2 on a MACS™ Dissociator. The cell suspension was then triturated through a 40 μm nylon mesh, and washed with PBS^-/-^ and total leukocyte numbers were determined. Single cell suspensions (<1x10^6^) from heart digests, spleens and peritoneal cells were incubated with a mixture of fluorophore-conjugated antibodies (Online Table 11) at 4°C for 30 minutes and then washed with PBS^-/-^. DAPI (1 μL/mL) was added to the samples for fluorescence associated cell sorting, immediately prior to acquisition.

**Flow cytometry and fluorescence associated cell sorting (FACS)**

Flow cytometric analysis was performed on an LSR II instrument (BD Biosciences) and analyzed using FlowJo software (Tree Star). Infarct zone CD45^+^CD11b^+^Ly6G^-^F4/80^+^ macrophages were sorted by FACS using a FACS ARIA II flow cytometer (BD Biosciences), see gating strategy in Online Figure 4. Results for the heart digests are expressed as cell number per infarct zone or remote zone, and total counts were calculated for spleens and the peripheral blood. Live single cells were gated for by excluding dead cells using DAPI, followed by singlet gates and subsequently by granularity and size. Fluorochrome conjugated antibodies were used to define cell populations of interest. Siglec-F and Ly6G surface markers were used to differentiate eosinophils (Siglec-F^+^Ly6G^int^) from neutrophils (Siglec-F^-^Ly6G^hi^).

**Detection of associated IL-4 by flow cytometry**

Release of IL-4 by heart or splenic leukocytes was measured using an IL-4 secretion assay (Miltenyi Biotec) according to the manufacturer's instructions. Briefly, single cell suspensions were stimulated for 2 hours with Cell Stimulation Cocktail (eBioscience). Subsequently cells were labelled with IL-4 Catch Reagent and incubated for further 45 minutes. Thereafter, cells were stained for cell surface markers as described above and analysed on a BD Fortessa flow cytometer (see Online Figure 7). As control unstimulated or cells incubated without Catch Reagent were used.

**Tissue Cytokine Assay**

Left ventricular infarct zone tissue was collected from ΔdblGATA and BALB/c mice, 4 days after induction of MI and snap frozen. Protein was extracted by bead mill homogenization of tissue in RIPA lysis and extraction buffer (Sigma-Aldrich) containing protease inhibitors (Sigma-Aldrich). Tissue cytokine amounts in the heart protein extracts were determined using the mouse Th2 Panel LEGENDplex assay (BioLegend), according to the manufactures instructions.
