## Supplementary figures and images for "Eosinophil deficiency promotes aberrant repair and adverse remodelling following acute myocardial infarction"

### Supplementary Figure 1

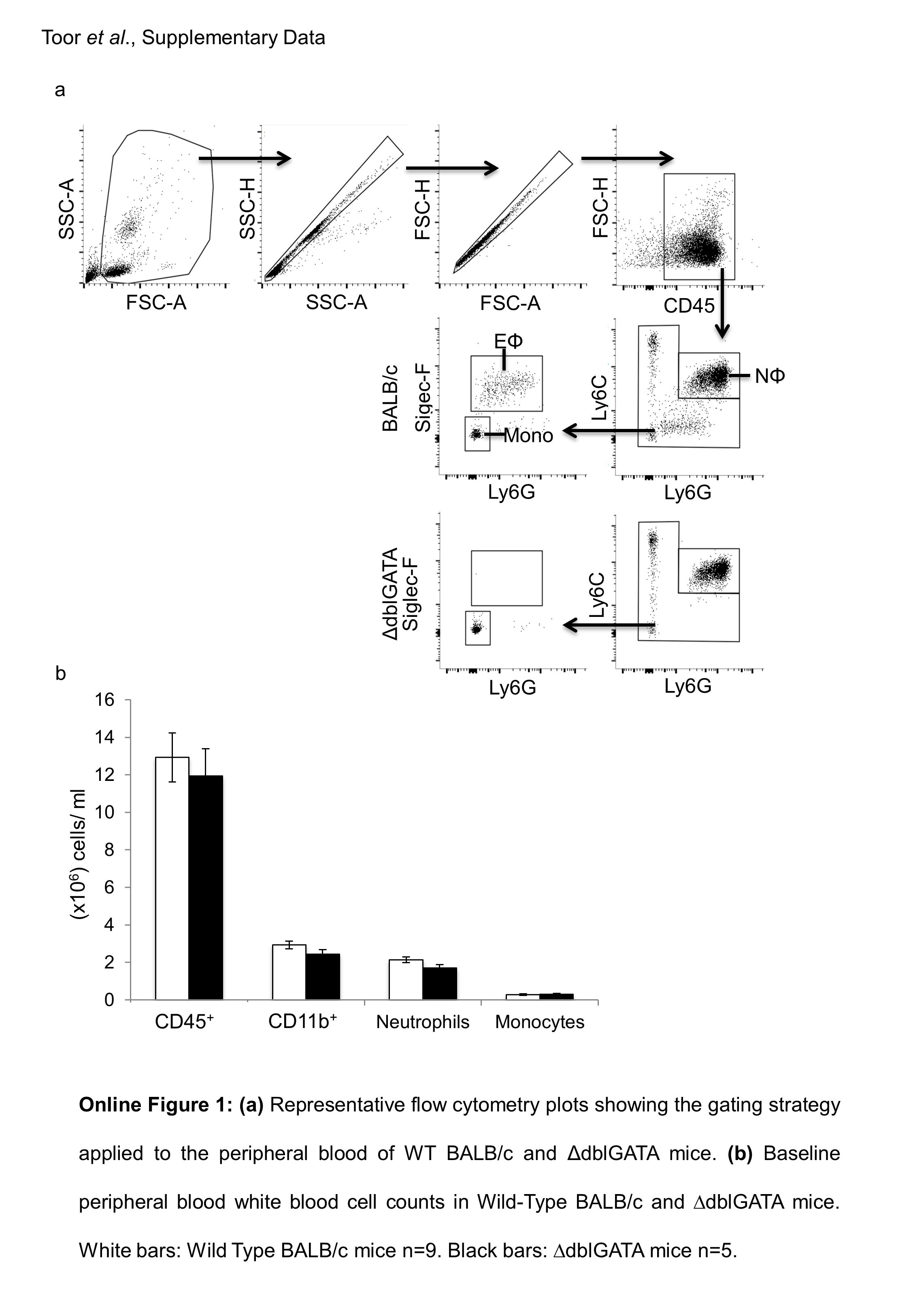
